# Simulating emotions: An active inference model of emotional state inference and emotion concept learning

**DOI:** 10.1101/640813

**Authors:** Ryan Smith, Thomas Parr, Karl J. Friston

**Author notes:** Corresponding Author Information: Ryan Smith, Laureate Institute for Brain Research, 6655 S Yale Ave, Tulsa, OK 74136, USA.

## Abstract

The ability to conceptualize and understand one’s own affective states and responses – or “emotional awareness” (EA) – is reduced in multiple psychiatric populations; it is also positively correlated with a range of adaptive cognitive and emotional traits. While a growing body of work has investigated the neurocognitive basis of EA, the neurocomputational processes underlying this ability have received limited attention. Here, we present a formal Active Inference (AI) model of emotion conceptualization that can simulate the neurocomputational (Bayesian) processes associated with learning about emotion concepts and inferring the emotions one is feeling in a given moment. We validate the model and inherent constructs by showing (i) it can successfully acquire a repertoire of emotion concepts in its “childhood”, as well as (ii) acquire new emotion concepts in synthetic “adulthood,” and (iii) that these learning processes depend on early experiences, environmental stability, and habitual patterns of selective attention. These results offer a proof of principle that cognitive-emotional processes can be modeled formally, and highlight the potential for both theoretical and empirical extensions of this line of research on emotion and emotional disorders.

---

The ability to conceptualize and understand one’s affective responses has become the topic of a growing body of empirical work (McRae, Reiman, Fort, Chen, & Lane, 2008; R. Smith, Alkozei, et al., 2017; R. Smith et al., 2015; R. Smith, Kaszniak, Katsanis, Lane, & Nielsen, 2019; R. Smith, Lane, Alkozei, et al., 2018; R. Smith, Lane, et al., 2017; R. Smith, Lane, Sanova, et al., 2018; R. Smith, Quinlan, et al., 2019; R. Smith, Sanova, Alkozei, Lane, & Killgore, 2018; Wright, Riedel, Sechrest, Lane, & Smith, 2017). This body of work has also given rise to theoretical models of its underlying cognitive and neural basis (Barrett, 2017; Kleckner et al., 2017; Lane, Weihs, Herring, Hishaw, & Smith, 2015; Panksepp, Lane, Solms, & Smith, 2017; R. Smith, Killgore, & Lane, 2018; R. Smith & Lane, 2015, 2016; Wilson-Mendenhall, Barrett, Simmons, & Barsalou, 2011). Attempts to operationalize this cognitive-emotional ability have led to a range of overlapping constructs, including trait emotional awareness (Lane & Schwartz, 1987), emotion differentiation or granularity (Kashdan, Barrett, & McKnight, 2015; Kashdan & Farmer, 2014), and alexithymia (Bagby, Parker, & Taylor, 1994a, 1994b).

This work is motivated to a large degree by the clinical relevance of emotion conceptualization abilities. In the literature on the construct of emotional awareness, for example, lower levels of conceptualization ability have been associated with several psychiatric disorders as well as poorer physical health (Baslet, Termini, & Herbener, 2009; Berthoz, Ouhayoun, & Parage, 2000; Bydlowski et al., 2005; Consoli et al., 2010; Donges et al., 2005; Frewen et al., 2008; Lackner, 2005; Levine, Marziali, & Hood, 1997; Moeller et al., 2014; A. Subic-Wrana, Beetz, Paulussen, Wiltnik, & Beutel, 2007; C. Subic-Wrana, Bruder, Thomas, Lane, & Köhle, 2005); conversely, higher ability levels have been associated with a range of adaptive emotion-related traits and abilities (Barchard & Hakstian, 2004; Bréjard, Bonnet, & Pedinielli, 2012; Ciarrochi, Caputi, & Mayer, 2003; Lane, Quinlan, Schwartz, Walker, & Zeitlin, 1990; Lane et al., 1996; Lane, Sechrest, Riedel, Shapiro, & Kaszniak, 2000). Multiple evidence-based psychotherapeutic modalities also aim to improve emotion understanding as a central part of psycho-education in psychotherapy (Barlow, Allen, & Choate, 2016; Hayes & Smith, 2005).

While there are a number of competing views on the nature of emotions, most (if not all) accept that emotion concepts must be acquired through experience. For example, “basic emotions” theories hold that emotion categories like sadness and fear each have distinct neural circuitry, but do not deny that knowledge about these emotions must be learned (Panksepp & Biven, 2012). Constructivist views instead hold that emotion categories do not have a 1-to-1 relationship to distinct neural circuitry, and that emotion concept acquisition is necessary for emotional experience (Barrett, 2017). While these views focus on understanding the nature of emotions themselves, we have recently proposed a neurocognitive model – termed the “three-process model” (TPM; (R. Smith, 2019; R. Smith, Kaszniak, et al., 2019; R. Smith, Killgore, et al., 2018)) of emotion episodes – with a primary focus on accounting for individual differences in emotional awareness. This model characterizes a range of emotion-related processes that could contribute to trait differences in both the learning and deployment of emotion concepts in order to understand one’s own affective responses (and in the subsequent use of these concepts to guide adaptive decision-making). The TPM distinguishes the following three broadly defined processes (see figure 1):

1. Affective response generation: a process in which somatovisceral and cognitive states are automatically modulated in response to an affective stimulus (whether real, remembered, or imagined) in a context-dependent manner, based on an (often implicit) appraisal of the significance of that stimulus for the survival and goal-achievement of the individual (i.e., predictions about the cognitive, metabolic, and behavioral demands of the situation).
2. Affective response representation: a process in which the somatovisceral component of an affective response is subsequently perceived via afferent sensory processing, and then conceptualized as a particular emotion (e.g., sadness, anger, etc.) in consideration of all other available sources of information (e.g., stimulus/context information, current thoughts/beliefs about the situation, etc.).
3. Conscious access: a process in which the representations of somatovisceral percepts and emotion concept representations may or may not enter and be held in working memory – constraining the use of this information in goal-directed decision-making (e.g., verbal reporting, selection of voluntary emotion regulation strategies, etc.).

**Figure 1.**
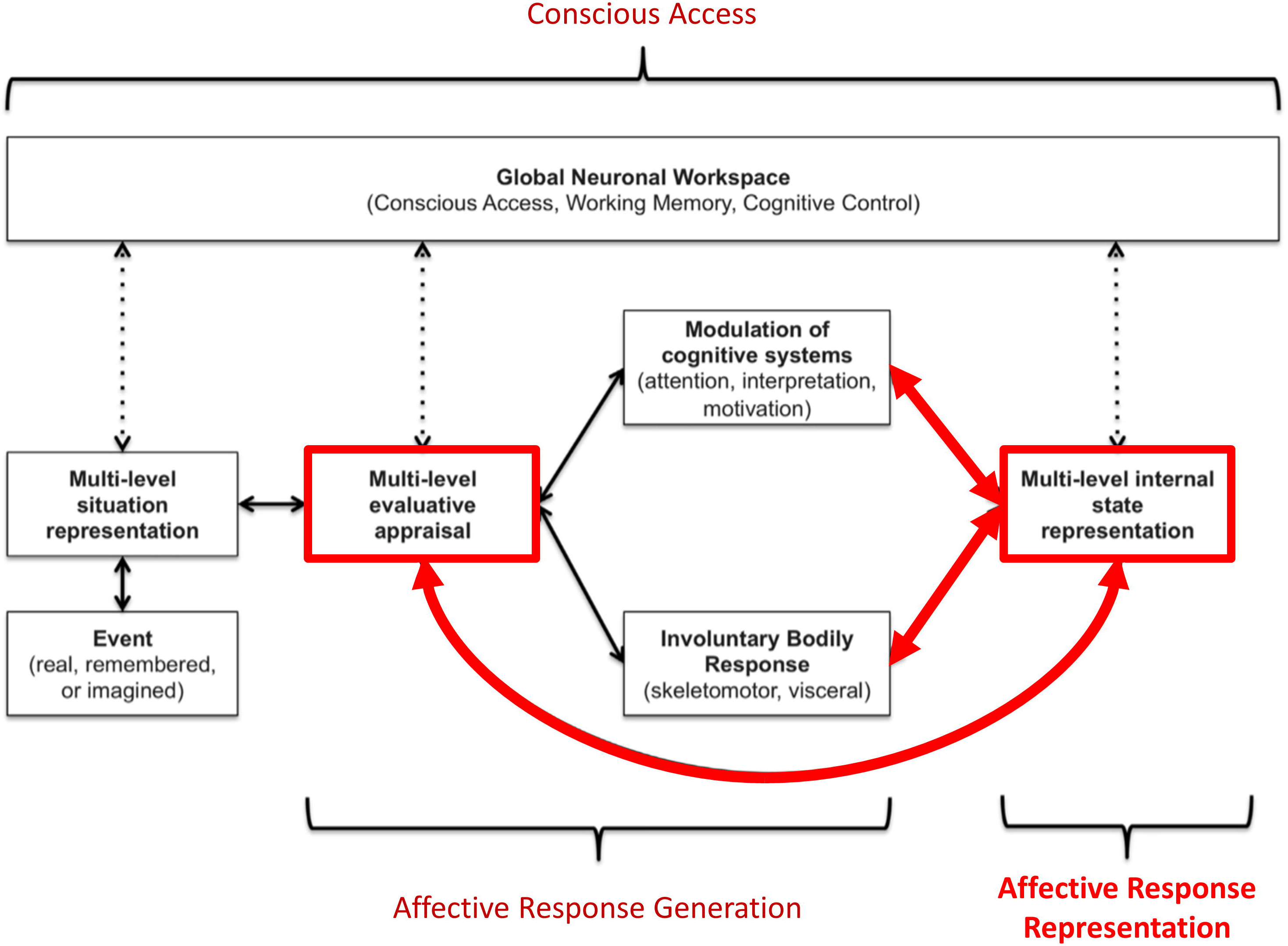
Graphical illustration of the three-process model of emotion episodes (R. Smith, 2019; R. Smith, Kaszniak, et al., 2019; R. Smith, Killgore, et al., 2018; R. Smith, Lane, Parr, & Friston, 2019). Here an event (either internal or external, and either real, remembered, or imagined) is represented in the brain at various levels of abstraction. This complex set of representations is evaluated in terms of its significance to the organism at both implicit (e.g., low-level conditioned or unconditioned responses) and explicit (e.g., needs, goals, values) levels, leading to predictions about the cognitive, metabolic, and behavioral demands of the situation. These predictions initiate an “affective response” with both peripheral and central components. This includes quick, involuntary autonomic and skeletomotor responses (e.g., heart rate changes, facial expression changes) as well as involuntary shifts in the direction of cognitive resources (e.g., attention and interpretation biases, action selection biases). This multicomponent affective response is represented centrally at a perceptual and conceptual level (e.g., valenced bodily sensations and interpretations of those responses as corresponding to particular emotions). Representations of events, evaluations, bodily sensations, emotion concepts (etc.) can then be attended to and gated into working memory (“conscious access”). If gated into and held in working memory, these representations can be integrated with explicit goals, reflected upon, and used to guide deliberative, goal-directed decision-making. The thick red arrows highlight the processes we explicitly model in this paper.

The TPM has also proposed a tentative mapping to the brain in terms of interactions between large-scale neural networks serving domain-general cognitive functions. Some support for this proposal has been found within recent neuroimaging studies (R. Smith, Alkozei, et al., 2017; R. Smith, Bajaj, et al., 2018; R. Smith, Lane, Alkozei, et al., 2018; R. Smith, Lane, et al., 2017; R. Smith, Lane, Sanova, et al., 2018; R. Smith, Sanova, et al., 2018). However, the neurocomputational implementation of these processes has not been thoroughly considered. The computational level of description offers the promise of providing more specific and mechanistic insights, which could potentially be exploited to inform and improve pharmacological and psychotherapeutic interventions. While previous theoretical work has applied active inference concepts to emotional phenomena (Barrett & Simmons, 2015; Clark, Watson, & Friston, 2018; Joffily & Coricelli, 2013; Seth, 2013; Seth & Friston, 2016; R. Smith, Thayer, Khalsa, & Lane, 2017; R. Smith, Weihs, Alkozei, Killgore, & Lane, 2019), no formal modeling of emotion concept learning has yet been performed. In this manuscript, we aim to take the first steps in constructing an explicit computational model of the acquisition and deployment of emotion concept knowledge (i.e., affective response representation) as described within the TPM (subsequent work will focus on affective response generation and conscious access processes; see (R. Smith, Lane, et al., 2019)). Specifically, we present a simple Active Inference model (Friston et al., 2016; Friston, FitzGerald, Rigoli, Schwartenbeck, & Pezzulo, 2017) of emotion conceptualization, formulated as a Markov Decision Process. We then outline some initial insights afforded by simulations using this model.

In what follows, we first provide a brief review of active inference. We will place a special emphasis on deep generative models that afford the capacity to explain multimodal (i.e., interoceptive, proprioceptive and exteroceptive) sensations that are characteristic of emotion concepts. We then introduce a particular model of emotion inference that is sufficiently nuanced to produce synthetic emotional processes but sufficiently simple to be understood from a ‘first principles’ account. We then establish the validity of this model using numerical analyses of emotion concept learning during (synthetic) neurodevelopment. We conclude with a brief discussion of the implications of this work; particularly for future applications.

## An Active Inference model of emotion conceptualization

### A primer on Active Inference

Active Inference (AI) starts from the assumption that the brain is an inference machine that approximates optimal probabilistic (Bayesian) belief updating across all biopsychological domains (e.g., perception, decision-making, motor control, etc.). AI postulates that the brain embodies an internal model of the world (including the body) that is “generative” in the sense that it is able to simulate the sensory data that it should receive if its model of the world is correct. This simulated (predicted) sensory data can be compared to actual observations, and deviations between predicted and observed sensations can then be used to update the model. On short timescales (e.g., a single trial in a perceptual decision-making task) this updating corresponds to perception, whereas on longer timescales it corresponds to learning (i.e., updating expectations about what will be observed on subsequent trials). One can see these processes as ensuring the generative model (embodied by the brain) remains an accurate model of the world (Conant & Ashbey, 1970).

Action (be it skeletomotor, visceromotor, or cognitive action) can be cast in similar terms. For example, actions can be chosen to resolve uncertainty about variables within a generative model (i.e., sampling from domains in which the model does not make precise predictions). This can prevent future deviations from predicted outcomes. In addition, the brain must continue to make certain predictions simply in order to survive. For example, if the brain did not in some sense continue to “expect” to observe certain amounts of food, water, shelter, social support, and a range of other quantities, then it would cease to exist (McKay & Dennett, 2009); as it would not pursue those behaviors leading to the realization of these expectations (c.f. ‘the optimism bias’ (Sharot, 2011)). Thus, there is a deep sense in which the brain must continually seek out observations that support – or are internally consistent with – its own continued existence. As a result, decision-making can be cast as a process in which the brain infers the sets of actions (policies) that would lead to observations most consistent with its own survival-related expectations (i.e., its “prior preferences”). Mathematically, this can be described as selecting policies that maximize a quantity called “Bayesian model evidence” – that is, the probability that sensory data would be observed under a given model. In other words, because the brain is itself a model of the world, action can be understood as a process by which the brain seeks out evidence for itself – sometimes known as self-evidencing (Hohwy, 2016).

In a real-world setting, directly computing model evidence becomes mathematically intractable. Thus, the brain must use some approximation. AI proposes that the brain instead computes a statistical quantity called free energy. Unlike model evidence, computing free energy is mathematically tractable. Crucially, this quantity provides a bound on model evidence, such that minimization of free energy is equivalent to maximizing model evidence. By extension, in decision-making an agent can evaluate the *expected* free energy of the alternative policies she could select – that is, the free energy of future trajectories under each policy (i.e., based on predicted future outcomes, given the future states that would be expected under each policy). Therefore, decision-making will be approximately (Bayes) optimal if it operates by inferring (and enacting) the policy that minimizes expected free energy – and thereby maximize evidence for the brain’s internal model. Interestingly, expected free energy can be decomposed into terms reflecting uncertainty and prior preferences, respectively. This decomposition explains why agents that minimizes expected free energy will first select exploratory policies that minimize uncertainty in a new environment (often called the “epistemic value” component of expected free energy). Once uncertainty is resolved, the agent then selects policies that exploit that environment to maximize her prior preferences (often called the “pragmatic value” component of expected free energy). The formal mathematical basis for AI has been detailed elsewhere (Friston, FitzGerald, et al., 2017), and the reader is referred there for a full mathematical treatment (also see figure 2 for some additional detail).

**Figure 2.**
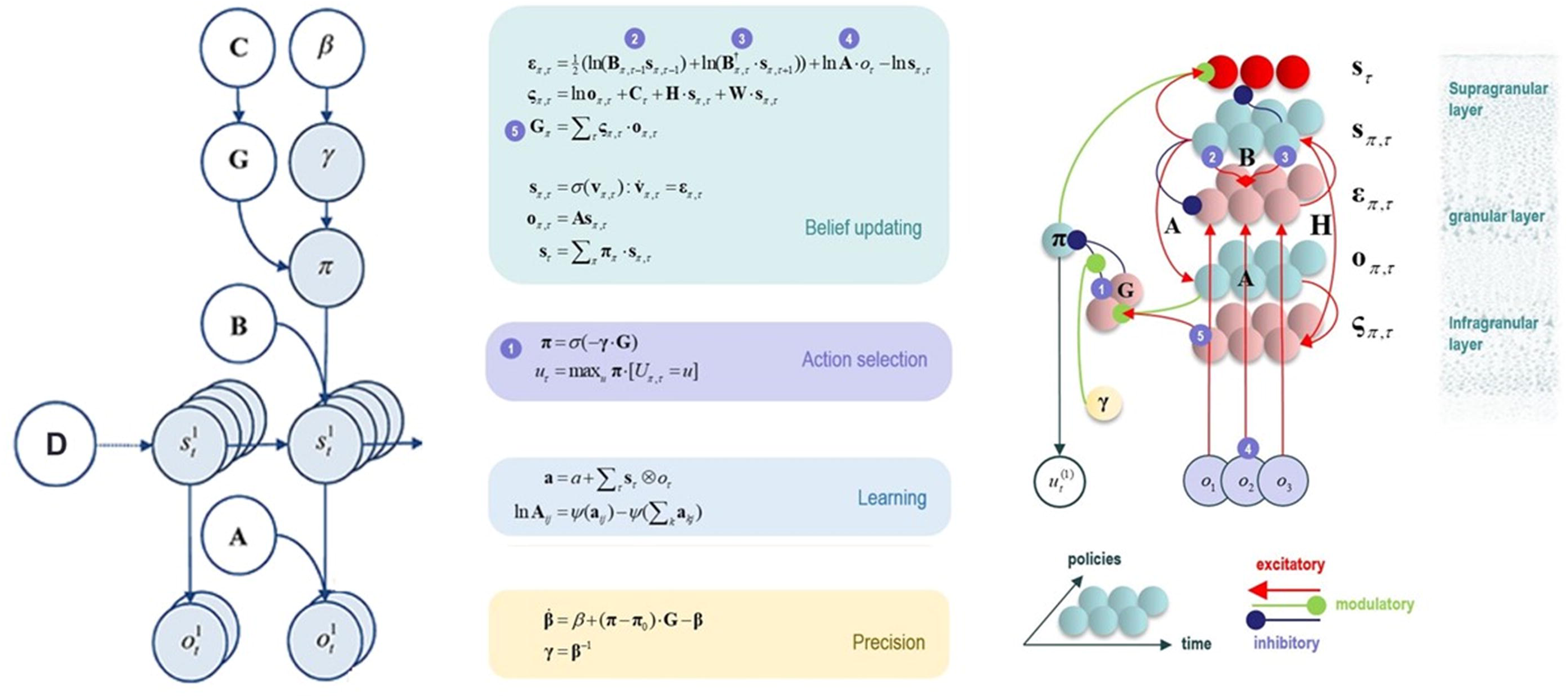
This figure illustrates the mathematical framework of active inference and associated neural process theory used in the simulations described in this paper. (Left) Illustration of the Markov decision process formulation of active inference. The generative model is here depicted graphically, such that arrows indicate dependencies between variables. Here observations (o) depend on hidden states (s), as specified by the A matrix, and those states depend on both previous states (as specified by the B matrix, or the initial states specified by the D matrix) and the policies (π) selected by the agent. The probability of selecting a particular policy in turn depends on the expected free energy (G) of each policy with respect to the prior preferences (C) of the agent. The degree to which expected free energy influences policy selection is also modulated by a prior policy precision parameter (γ), which is in turn dependent on beta (*β*) –where higher values of beta promote more randomness in policy selection (i.e., less influence of the differences in expected free energy across policies). (Middle/Right) The differential equations in the middle panel approximate Bayesian belief updating within the graphical model depicted on the left via a gradient descent on free energy (F). The right panel also illustrates the proposed neural basis by which neurons making up cortical columns could implement these equations. The equations have been expressed in terms of two types of prediction errors. State prediction errors (ε) signal the difference between the (logarithms of) expected states (s) under each policy and time point—and the corresponding predictions based upon outcomes/observations (A matrix) and the (preceding and subsequent) hidden states (B matrix, and, although not written, the D matrix for the initial hidden states at the first time point). These represent prior and likelihood terms respectively – also marked as messages 2, 3, and 4, which are depicted as being passed between neural populations (colored balls) via particular synaptic connections in the right panel (note: the dot notation indicates transposed matrix multiplication within the update equations for prediction errors). These (prediction error) signals drive depolarization (v) in those neurons encoding hidden states (s), where the probability distribution over hidden states is then obtained via a softmax (normalized exponential) function (σ). Outcome prediction errors (ς) instead signal the difference between the (logarithms of) expected observations (o) and those predicted under prior preferences (C). This term additionally considers the expected ambiguity or conditional entropy (H) between states and outcomes as well as a novelty term (W) reflecting the degree to which beliefs about how states generate outcomes would change upon observing different possible state-outcome mappings (computed from the A matrix). This prediction error is weighted by the expected observations to evaluate the expected free energy (G) for each policy (π), conveyed via message 5. These policy-specific free energies are then integrated to give the policy expectations via a softmax function, conveyed through message 1. Actions at each time point (u) are then chosen out of the possible actions under each policy (U) weighted by the value (negative expected free energy) of each policy. In our simulations, the model learned associations between hidden states and observations (A) via a process in which counts were accumulated (a) reflecting the number of times the agent observed a particular outcome when she believed that she occupied each possible hidden state. Although not displayed explicitly, learning prior expectations over initial hidden states (D) is similarly accomplished via accumulation of concentration parameters (d). These prior expectations reflect counts of how many times the agent believes it previously occupied each possible initial state. Concentration parameters are converted into expected log probabilities using digamma functions (*ψ*). As already stated, the right panel illustrates a possible neural implementation of the update equations in the middle panel. In this implementation, probability estimates have been associated with neuronal populations that are arranged to reproduce known intrinsic (within cortical area) connections. Red connections are excitatory, blue connections are inhibitory, and green connections are modulatory (i.e., involve a multiplication or weighting). These connections mediate the message passing associated with the equations in the middle panel. Cyan units correspond to expectations about hidden states and (future) outcomes under each policy, while red states indicate their Bayesian model averages (i.e., a “best guess” based on the average of the probability estimates for the states and outcomes across policies, weighted by the probability estimates for their associated policies). Pink units correspond to (state and outcome) prediction errors that are averaged to evaluate expected free energy and subsequent policy expectations (in the lower part of the network). This (neural) network formulation of belief updating means that connection strengths correspond to the parameters of the generative model described in the text. Learning then corresponds to changes in the synaptic connection strengths. Only exemplar connections are shown to avoid visual clutter. Furthermore, we have just shown neuronal populations encoding hidden states under two policies over three time points (i.e., two transitions), whereas in the task described in this paper there are greater number of allowable policies. For more information regarding the mathematics and processes illustrated in this figure, see (Friston et al., 2016; Friston, FitzGerald, et al., 2017; Friston, Lin, et al., 2017; Friston, Parr, & de Vries, 2017).

When a generative model is formulated as a partially observable Markov decision process, active inference takes a particular form. Specifically, specifying a generative model in this context requires specifying the allowable policies, hidden states of the world (that the brain cannot directly observe but must infer), and observable outcomes, as well as a number of matrices that define the probabilistic relationships between these quantities (see figure 2). The ‘A’ matrix specifies which outcomes are generated by each combination of hidden states (i.e., a likelihood mapping indicating the probability that a particular set of outcomes would be observed given a particular hidden state). The ‘B’ matrix encodes state transitions, specifying the probability that one hidden state will evolve into another over time. Some of these transitions are controlled by the agent, according to the policy that has been selected. The ‘D’ matrix encodes prior expectations about the initial hidden state of the world. The ‘E’ matrix encodes prior expectations about which policies will be chosen (e.g., frequently repeated habitual behaviors will have higher prior expectation values). Finally, the ‘C’ matrix encodes prior preferences over outcomes. Outcomes and hidden states are generally factorized into multiple outcome *modalities* and hidden state *factors*. This means that the likelihood mapping (the ‘A’ matrix) plays an important role in modeling the interactions among different hidden states at each level of a hierarchical model when generating the outcomes at the level below. One can think of each factor and modality as an independent group of competing states or observations within a given category. For example, one hidden state factor could be “birds,” which includes competing interpretations of sensory input as corresponding to either hawks, parrots, or pigeons, whereas a separate factor could be “location,” with competing representations of where a bird is in the sky. Similarly, one outcome modality could be size (e.g., is the bird big or small?) whereas another could be color (is the bird black, white, green, etc.?).

As shown in the middle and right panels of figure 2, active inference is also equipped with a neural process theory – a proposed manner in which neuronal circuits and their dynamics can invert generative models via a set of linked update equations that minimize prediction errors. In this neuronal implementation, the probability of neuronal firing in specific populations is associated with the expected probability of a state, whereas postsynaptic membrane potentials are associated with the logarithm of this probability. A softmax function acts as an activation function – transforming membrane potentials into firing rates. With this setup, postsynaptic depolarizations (driven by ascending signals) can be understood as prediction errors (free energy gradients) about hidden states – arising from linear mixtures in the firing rates of other neural populations. These prediction errors (postsynaptic currents) in turn drive membrane potential changes (and resulting firing rates). When predictions errors are minimized, postsynaptic influences no longer drive changes in activity (depolarizations and firing rates), corresponding to minimum free energy.

Via similar dynamics, predictions errors about outcomes (i.e., the deviation between preferred outcomes and those predicted under each policy) can also be computed and integrated (i.e., averaged) to evaluate the expected free energy (value) of each policy (i.e., underwriting selection of the policy that best minimizes these prediction errors). Dopamine dynamics also modulate policy selection, by encoding estimates of the expected uncertainty over policies – where greater expected uncertainty promotes less deterministic policy selection. Phasic dopamine responses correspond to updates in expected uncertainty over policies – which occur when there is a prediction error about expected free energy; that is, when there is a difference between the expected free energy of policies before and after a new observation.

Finally, and most centrally for the simulations we report below, learning in this theory corresponds to a form of synaptic plasticity remarkably similar to Hebbian coincidence-based learning mechanisms associated with empirically observed synaptic long-term potentiation and depression (LTP and LTD) processes (Brown, Zhao, & Leung, 2010). Here one can think of the strength of each synaptic connection as a parameter in one of the matrices described above. For example, the strength of one synapse could encode the amount of evidence a given observation provides for a given hidden state (i.e., an entry in the ‘A’ matrix), whereas another synapse could encode the probability of a state at a later time given a state at an earlier time (i.e., an entry in the ‘B’ matrix). Mathematically, the synaptic strengths correspond to Dirichlet parameters that increase in value in response to new observations. One can think of this process as adding counts to each matrix entry based on coincidences in pre- and post-synaptic activity. For example, if beliefs favor one hidden state, and this co-occurs with a specific observation, then the strength of the value in the ‘A’ matrix encoding the relationship between that state and that observation will increase. Counts also increase in similar fashion in the ‘D’ matrix encoding prior beliefs about initial states (whenever a given hidden state is observed at the start of a trial) as well as in the ‘B’ matrix encoding beliefs about transition probabilities (whenever one specific state is followed by another). For a more detailed discussion, please see the legend for figure 2 and associated references.

In what follows, we describe how this type of generative model was specified to perform emotional state inference and emotion concept learning. We also present simulated neural responses based on the neural process theory described above.

### A model of emotion inference and concept learning

In this paper we focus on the second process in the TPM – affective response representation – in which a multifaceted affective response is generated and the ensuing (exteroceptive, proprioceptive and interoceptive) outcomes are used to infer or represent the current emotional state. The basic idea is to equip the generative model with a space of emotion concepts (i.e., latent or hidden states) that generate the interoceptive, exteroceptive and proprioceptive consequences (at various levels of abstraction) of being in a particular emotional state. Inference under this model then corresponds to inferring that one of several possible emotion concepts is the best explanation for the data at hand (e.g., “my unpleasant feeling of increased heart rate and urge to run away must indicate that I am afraid to give this speech”). Crucially, to endow emotion concept inference with a form of mental action (Limanowski & Friston, 2018; Metzinger, 2017), we also included a state factor corresponding to *selective attention*. Transitions between attentional states were under control of the agent (i.e., ‘B’ matrices were specified for all possible transitions between these states). The ‘A’ matrix mapping emotion concepts to lower-level observations differed in each attentional state, such that precise information about each type of lower-level information was only available in one attentional state (e.g., the agent needed to transition into the “attention to valence” state to gain precise information whether she was feeling pleasant or unpleasant, and so forth; see figure 3C).

The incorporation of selective attention in emotional state inference and learning within our model was motivated by several factors. First, multiple psychotherapeutic modalities improve clients’ understanding of their own emotions in just this way; that is, by having them selectively attend to and record the contexts, bodily sensations, thoughts, action tendencies, and behaviours during emotion-episodes (e.g., (Barlow et al., 2016; Hayes & Smith, 2005)). Second, low emotional awareness has been linked to biased attention in some clinical contexts (Lane, Anderson, & Smith, 2018). Third, related personality factors (e.g., biases toward “externally oriented thinking”) are included in leading self-report measures of the related construct of alexithymia (Parker, Taylor, & Bagby, 2003). Finally, emotion learning in childhood appears to involve parent-child interactions in which parents draw attention to (and label) bodily feelings and behaviors during a child’s affective responses (e.g., see work on attunement, social referencing, and related aspects of emotional development (Licata, Kristen, & Sodian, 2016; Mumme, Fernald, & Herrera, 1996; R. Smith, Killgore, et al., 2018)) – and the lack of such interactions hinders emotion learning (and mental state learning more generally; (Colvert et al., 2008)).

In our model, we used relatively high level ‘outcomes’ (i.e., themselves standing in for lower-level representations) to summarise the products of belief updating at lower levels of a hierarchical model. These outcomes were domain-specific, covering interoceptive, proprioceptive and exteroceptive modalities. A full hierarchical model would consider lower levels, unpacked in terms of sensory modalities; however, the current model, comprising just two levels, is sufficient for our purposes. The bottom portion of Figure 3A (in grey) acknowledges the broad form that these lower-level outcomes would be expected to take. The full three-process model would also contain a higher level corresponding to conscious accessibility (for an explicit model and simulations of this higher level, see (R. Smith, Lane, et al., 2019)). This is indicated by the grey arrows at the top of Figure 3A.

**Figure 3.**
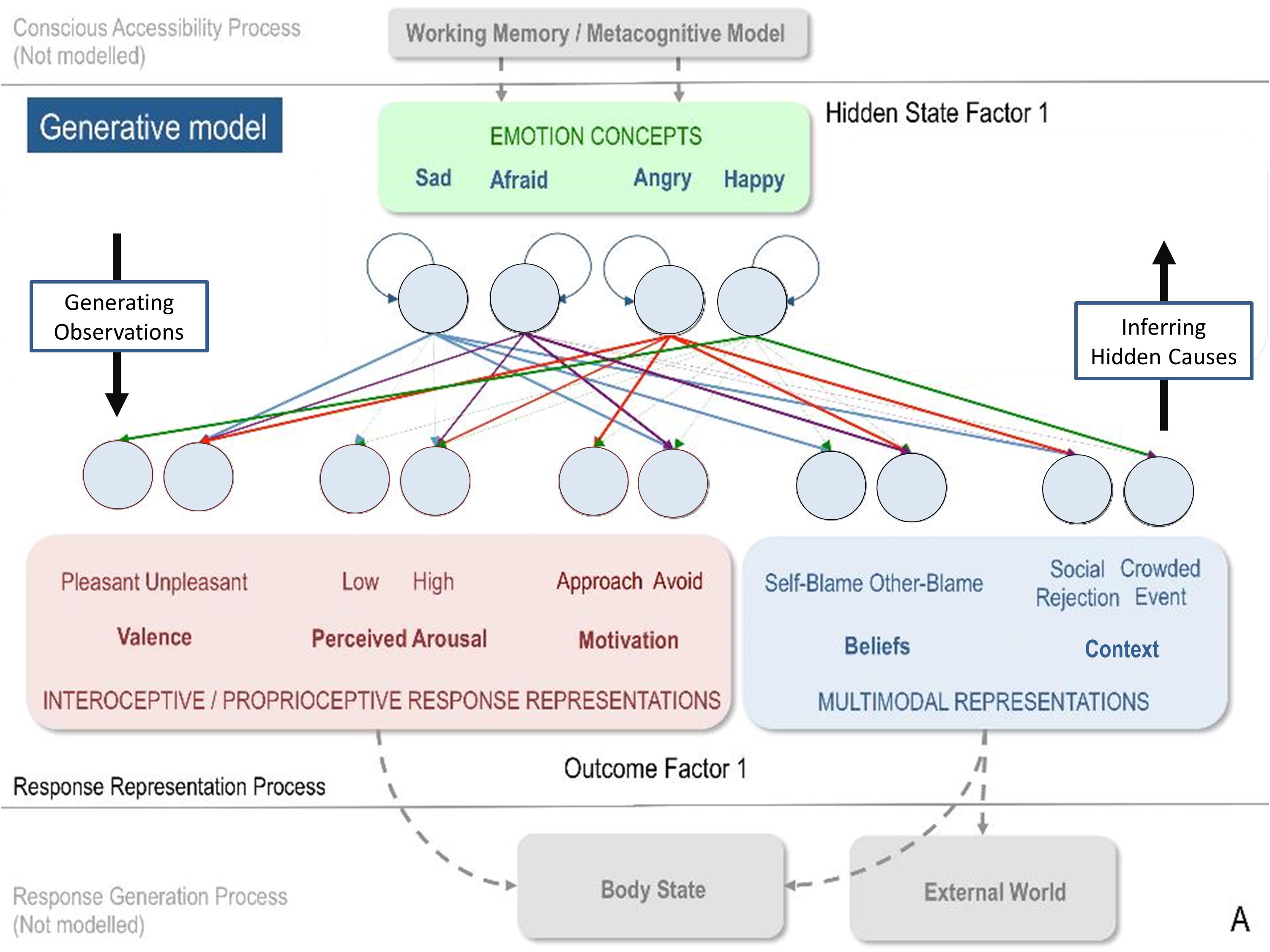
(A) Displays the levels of hidden state factor 1 (emotion concepts) and their mapping to different lower-level representational contents (here modelled as outcomes). Each emotion concept generated different outcome patterns (see text for details), although some were more specific than others (e.g., HAPPY generated high and low arousal equally; denoted by thin dotted connections). The A-matrix encoding these mappings is shown in figure 3C. The B-matrix, also shown in figure 3C, was an identity mapping between emotion states, such that emotions were stable within trials. The precision of this matrix (i.e., implicit beliefs about emotional stability) could be adjusted via passing this matrix through a softmax function with different temperature parameter values. This model simulates the affective response representation process within the three-process model of emotion episodes (R. Smith, 2019; R. Smith, Kaszniak, et al., 2019; R. Smith, Killgore, et al., 2018). Black arrows on the right and left indicate the direction by which hidden causes generate observations (the generative process) and the direction of inference in which observations are used to infer their hidden causes (emotion concepts) using the agent’s generative model. The grey arrows/boxes at the bottom and top of the figure denote other processes within the three-process model (i.e., affective response generation and conscious accessibility) that are not explicitly modeled in the current work (for simulations of the conscious accessibility process, see (R. Smith, Lane, et al., 2019)).

Crucially, as mentioned above, attentional focus was treated as a (mental) action that determines the outcome modality or domain to which attention was selectively allocated. Effectively, the agent had to decide which lower-level representations to selectively attend to (i.e., which sequential attention policy to select) in order to figure out what emotional state she was in. Mathematically, this was implemented via interactions in the likelihood mapping – such that being in a particular attentional state selected one and only one precise mapping between the emotional state factor and the outcome information in question (see figure 3C). Formally, this implementation of mental action or attentional focus is exactly the same used to model the exploration of a visual scene using overt eye movements (Mirza, Adams, Mathys, & Friston, 2016). However, on our interpretation, this epistemic foraging was entirely covert; hence mental action (c.f., the premotor theory of attention; (Posner, 2016; Rizzolatti, Riggio, Dascola, & Umiltá, 1987; D. Smith & Schenk, 2012)).

Figure 3 illustrates the resulting model. The first hidden state factor was a space of emotion concepts (SAD, AFRAID, ANGRY, and HAPPY). The second hidden state factor was attentional focus, and the ‘B’ matrix for this second factor allowed state transitions to be controlled by the agent. The agent could choose to attend to three sources of bodily (interoceptive/proprioceptive) information, corresponding to affective valence (pleasant or unpleasant sensations), autonomic arousal (e.g., high or low heart rate), and motivated proprioceptive action tendencies (approach or avoid). The agent could also attend to two sources of exteroceptive information, including the perceived situation (involving social rejection or a crowded event) and subsequent beliefs about responsibility (attributing agency/blame to self or another). These different sources of information are based on a large literature within emotion research, indicating that they are jointly predictive of self-reported emotions and/or are important factors in affective processing (Barrett, 2017; Barrett, Mesquita, & Gendron, 2011; Harmon-Jones, Gable, & Peterson, 2010; Lindquist & Barrett, 2008; Russell, 2003; Scherer, 2009; Siemer, Mauss, & Gross, 2007).

Our choice of including valence in particular reflects the fact that our model deals with high levels of hierarchical processing (this choice also enables us to connect more fluently with current literature on emotion concept categories). In this paper, we are using labels like ‘unpleasant’ as pre-emotional constructs. In other words, although affective in nature, we take concepts like ‘unpleasant’ as contributing to elaborated emotional constructs during inference. Technically, valenced states provide evidence for – or are a consequence of – emotional state inference at a higher level (e.g., pleasant sensations provide evidence that one is feeling a positive emotion like excitement, joy, or contentment, whereas unpleasant sensations provide evidence that one may be feeling a negative emotion such as as sadness, fear, or anger). Based on previous work (Clark et al., 2018; Joffily & Coricelli, 2013), we might expect valence to correspond to changes in the precision/confidence associated with lower-level visceromotor and skeletomotor policy selection, or to related internal estimates that can act as indicators of success in uncertainty resolution; see (Clark et al., 2018; de Berker et al., 2016; Joffily & Coricelli, 2013; Peters, McEwen, & Friston, 2017)). Put another way, feeling good may correspond to high confidence in one’s model of how to act, whereas feeling bad may reflect the opposite. Explicitly modelling these lower-level dynamics in a deep temporal model will be the focus of future work.

**Figure 3.**
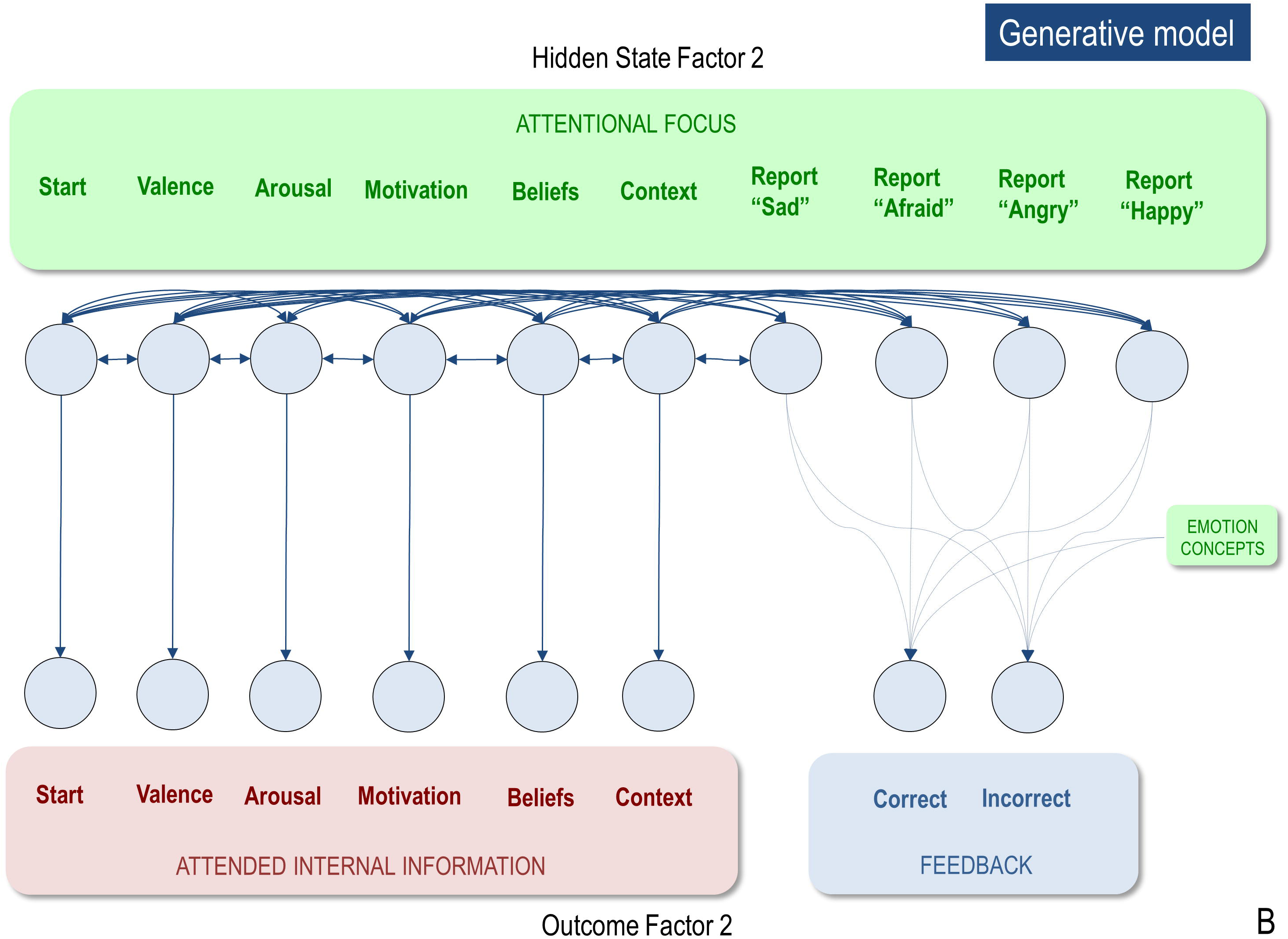
(B) Displays the levels of hidden state factor 2 (focus of attention) and its mapping to representational outcomes. Each focus of attention mapped deterministically (the A-matrix was a fully precise identity matrix) to a “location” (i.e., an internal source of information) at which different representational outcomes could be observed. Multiple B-matrices, depicted in figure 3C, provided controllable transitions (i.e., actions) such that the agent could choose to shift her attention from one internal representation to another to facilitate inference. The agent always began a trial in the “start” attentional state, which provided no informative observations. The final attentional shift in the trial was toward a (proprioceptive) motor response to report an emotion (i.e., at whatever point in the trial the agent became sufficiently confident, at which point the state could not change until the end of the trial), which was either correct or incorrect. The agent preferred (expected) to be correct and not to be incorrect. Because policies (i.e., sequences of implicit attentional shifts and subsequent explicit reports) were selected to minimize expected free energy, emotional state inference under this model entails a sampling of salient representational outcomes and subsequent report – under the prior preference that the report would elicit the outcome of “correct” social feedback. In other words, policy selection was initially dominated by the epistemic value part of expected free energy (driving the agent to gather information about her emotional state); then, as certainty increased, the pragmatic value part of expected free energy gradually began to dominate (driving the agent to report her emotional state).

In the simulations we report here, there were 6 time points in each epoch or trial of emotion inference. At the first time point, the agent always began in an uninformative initial state of attentional set (the “start” state). The agent’s task was to choose what to attend to, and in which order, to infer her most likely emotional state. When she became sufficiently confident, she could choose to respond (i.e., reporting that she felt sad, afraid, angry, or happy). In these simulations the agent selected “shallow” one-step policies, such that she could choose what to attend to next – to gain the most information. Given the number of time points, the agent could choose to attend to up to four of the five possible sources of outcome information before reporting her beliefs about her emotional state. The ‘A’ matrix mapping attentional focus to attended outcomes was an identity matrix, such that the agent always knew which lower-level information she was currently attending to. This may be thought of as analogous to the proprioceptive feedback consequent on a motor action.

**Figure 3.**
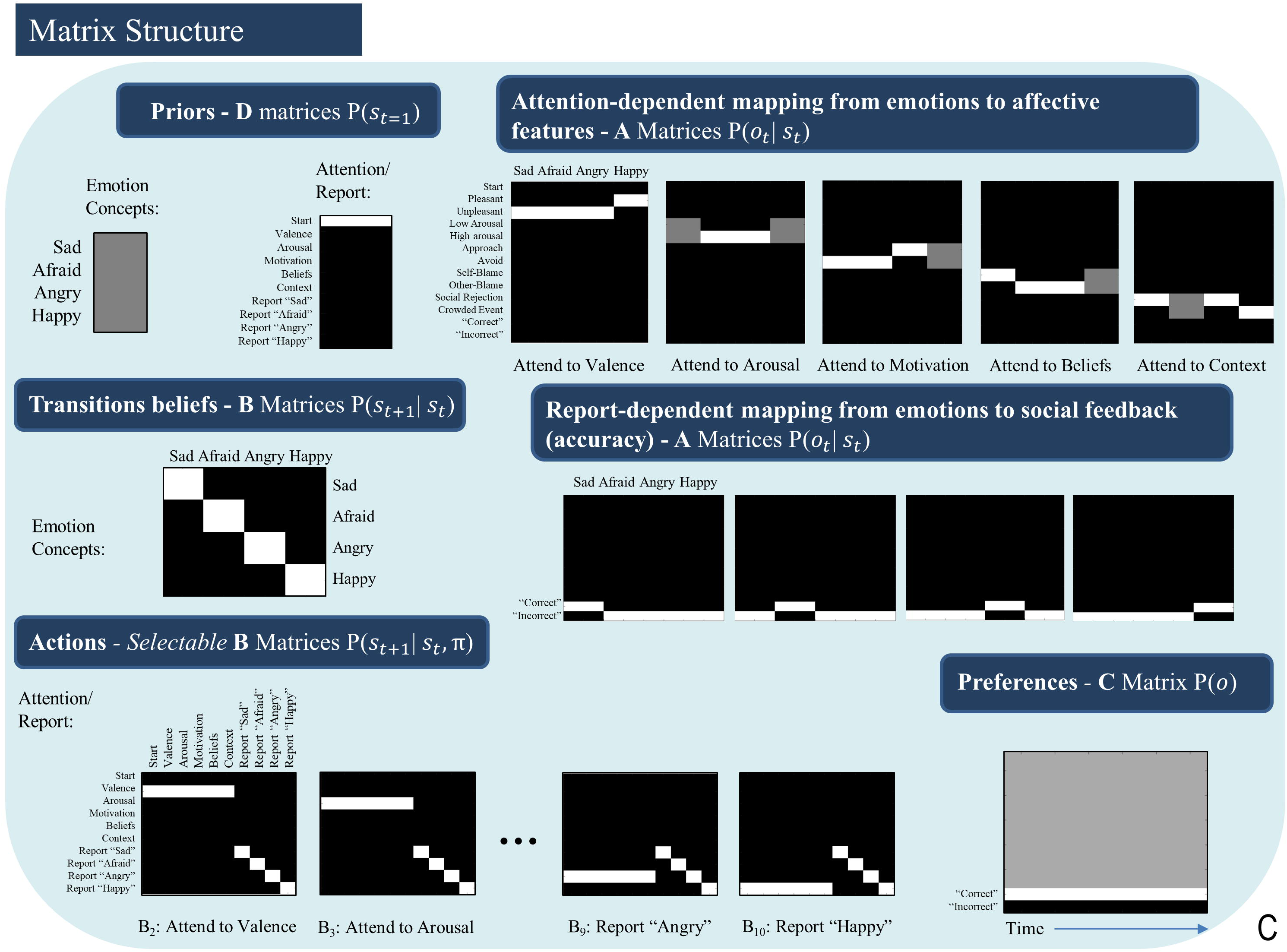
(C) Displays select matrices defining the generative model (lighter colors indicate higher probabilities). ‘D’ matrices indicate a flat distribution over initial emotional states and a strong belief that the attentional state will begin in the “start” state. ‘A’ matrices indicate the observations (rows) that would be generated by each emotional state (columns) depending on the current state of attention, as well as the social feedback (accuracy information) that would be generated under each emotional state depending on chosen self-reports. ‘B’ matrices indicate that emotional states are stable within a trial (i.e., states transition only to themselves such that the state transition matrix is an identity matrix) and that the agent can choose to shift attention to each modality of lower-level information (i.e., by selecting the transitions encoded by ‘B’ matrices 2-6), or report her emotional state (i.e., by selecting the transitions entailed by ‘B’ matrices 7-10), at which point she could no longer leave that state. Only the first and last two possible attentional shifting actions (‘B’ matrices) are shown due to space constraints (note: the first ‘B’ matrix for this factor corresponds to remaining in the starting state and is not shown); ‘B’ matrices 4-8 take identical form, but with the row vector [1 1 1 1 1 1] spanning the first 6 columns progressively shifted downward with each matrix, indicating the ability to shift from each attentional state to the attentional state corresponding to that row. The ‘C’ matrix indicates a preference for “correct” social feedback and an aversion to “incorrect” social feedback.

The ‘B’ matrix for hidden emotional states was also an identity matrix, reflecting the belief that emotional states are stable within a trial (i.e., if you start out feeling sad, then you will remain sad throughout the trial). This sort of probability transition matrix in the generative model allows evidence to be accumulated for one state or another over time; here, the emotion concept that provides the best explanation for actively attended evidence in the outcome modalities. The ‘A’ matrix – mapping emotion concepts to outcomes – was constructed such that certain outcome combinations were more consistent with certain emotional states than others: SAD was probabilistically associated with unpleasant valence, either low or high arousal (e.g., lying in bed lethargically vs. intensely crying), avoidance motivation, social rejection, and self-attribution (i.e., self-blame). AFRAID generated unpleasant valence, high arousal, avoidance, other-blame (c.f., fear often being associated with its perceived external cause), and either social rejection or a crowded event (e.g., fear of a life without friends vs. panic in crowded spaces). ANGRY generated unpleasant valence, high arousal, approach, social rejection, and other-blame outcomes. Finally, HAPPY generated pleasant valence, either low or high arousal, either approach or avoidance (e.g., feeling excited to wake up and go to work vs. feeling content in bed and not wanting to go to work), and a crowded event (e.g., having fun at a concert). Because HAPPY does not have strong conceptual links to blame, we defined a flat mapping between HAPPY and blame, such that either type of blame provided no evidence for or against being happy. Although this mapping from emotional states to outcomes has some face validity, it should not be taken too seriously. It was chosen primarily to capture the ambiguous and overlapping correlates of emotion concepts, and to highlight why adaptive emotional state inference and emotion concept learning can represent difficult problems.

If the ‘A’ matrix encoding state-outcome relationships was completely precise (i.e., if the contingencies above were deterministic as opposed to probabilistic), sufficient information could be gathered through (at most) three attentional shifts; but this becomes more difficult when probabilistic mappings are imprecise (i.e., as they more plausibly are in the real world). Figure 4 illustrates this by showing how the synthetic subjects confidence about her state decreases as the precision of the mapping between emotional states and outcomes decreases (we measured confidence here in terms of the accuracy of responding in relation to the same setup with infinite precision). Changes in precision were implemented via a temperature parameter of a softmax function applied to a fully precise version of likelihood mappings between emotion concepts and the 5 types of lower-level information that the agent could attend to (where a higher value indicates higher precision). For a more technical account of this type of manipulation, please see (Parr & Friston, 2017a).

Figure 4 additionally demonstrates how reporting confidence decreases with decreasing precision of the ‘B’ matrix encoding emotional state transitions, where low precision corresponds to the belief that emotional states are unstable over time. Interestingly, these results suggest that expectations about emotional instability would reduce the ability to understand or infer one’s own emotions. From a Bayesian perspective, this result is very sensible: if we are unable to use past beliefs to contextualise the present, it is much harder to accumulate evidence in favour of one hypothesis about emotional state relative to another. Under moderate levels of precision, our numerical analysis demonstrates that the model can conceptualize the multimodal affective responses it perceives with high accuracy.

**Figure 4.**
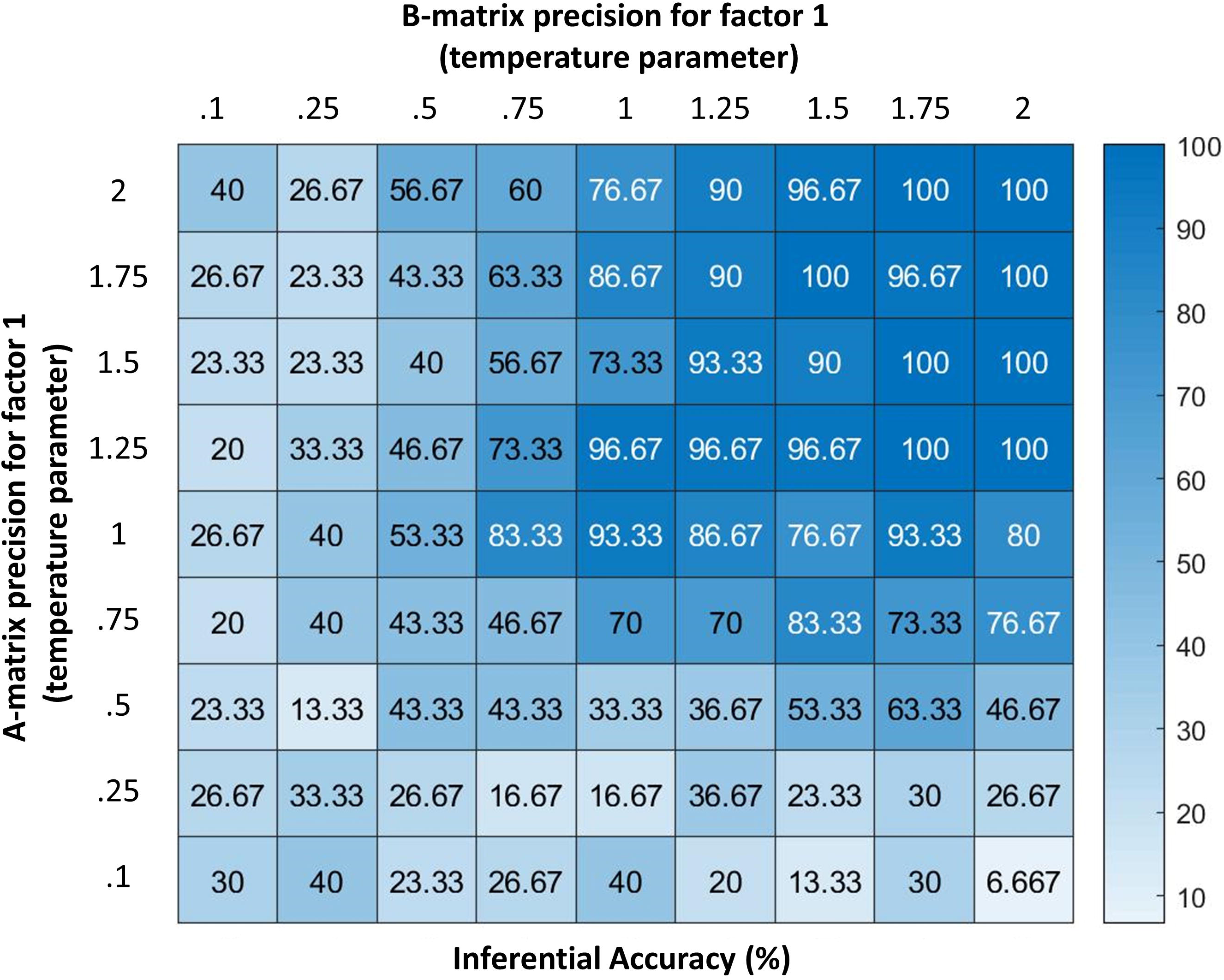
Displays the accuracy of the model (percentage of correct inferences over 30 trials) under different levels of precision for two parameters (denoted by temperature values for a softmax function controlling the specificity of the A and B matrices for hidden state factor 1; higher values indicate higher precision). As can be seen, the model performs with high accuracy at moderate levels of precision. However, its ability to infer its own emotions becomes very poor if the precision of either matrix becomes highly imprecise. Accuracy here is defined in relation to the response obtained from an agent with infinite precision – and can be taken as a behavioural measure of the quality of belief updating about emotional states. These results illustrate how emotion concepts could be successfully inferred despite variability in lower-level observations (e.g., contexts, arousal levels), as would be expected under constructivist theories of emotion (Barrett, 2017); however, they also demonstrate limits in variability, beyond which self-focused emotion recognition would begin to fail.

Figure 5 illustrates a range of simulation results from an example trial under moderately high levels of ‘A’ and ‘B’ matrix precision (temperature parameter = 2 for each). The upper left plot shows the sequence of (inferred) attentional shifts (note: darker colours indicate higher probability beliefs of the agent, and cyan dots indicate the true states). In this trial, the agent chose a policy in which it attended to valence (observing “unpleasant”), then beliefs (observing “other-blame”), then action (observing “approach”), at which time she became sufficiently confident and chose to report that she was angry. The lower left plot displays the agent’s *posterior* beliefs at the end of the trial about her emotional state at each timepoint in the trial, in this case inferring that she had been (and still was) angry. Note that this reflects retrospective inference, and not the agent’s beliefs at each timepoint. The lower right and upper right plots display simulated neural responses (based on the neural process theory that accompanies this form of active inference; (Friston, FitzGerald, et al., 2017)), in terms of single-neuron firing rates (raster plots) and local field potentials, respectively. The simulated firing rates in the lower right plot illustrate that the agent’s confidence that she was angry increased gradually with each new observation.

**Figure 5.**
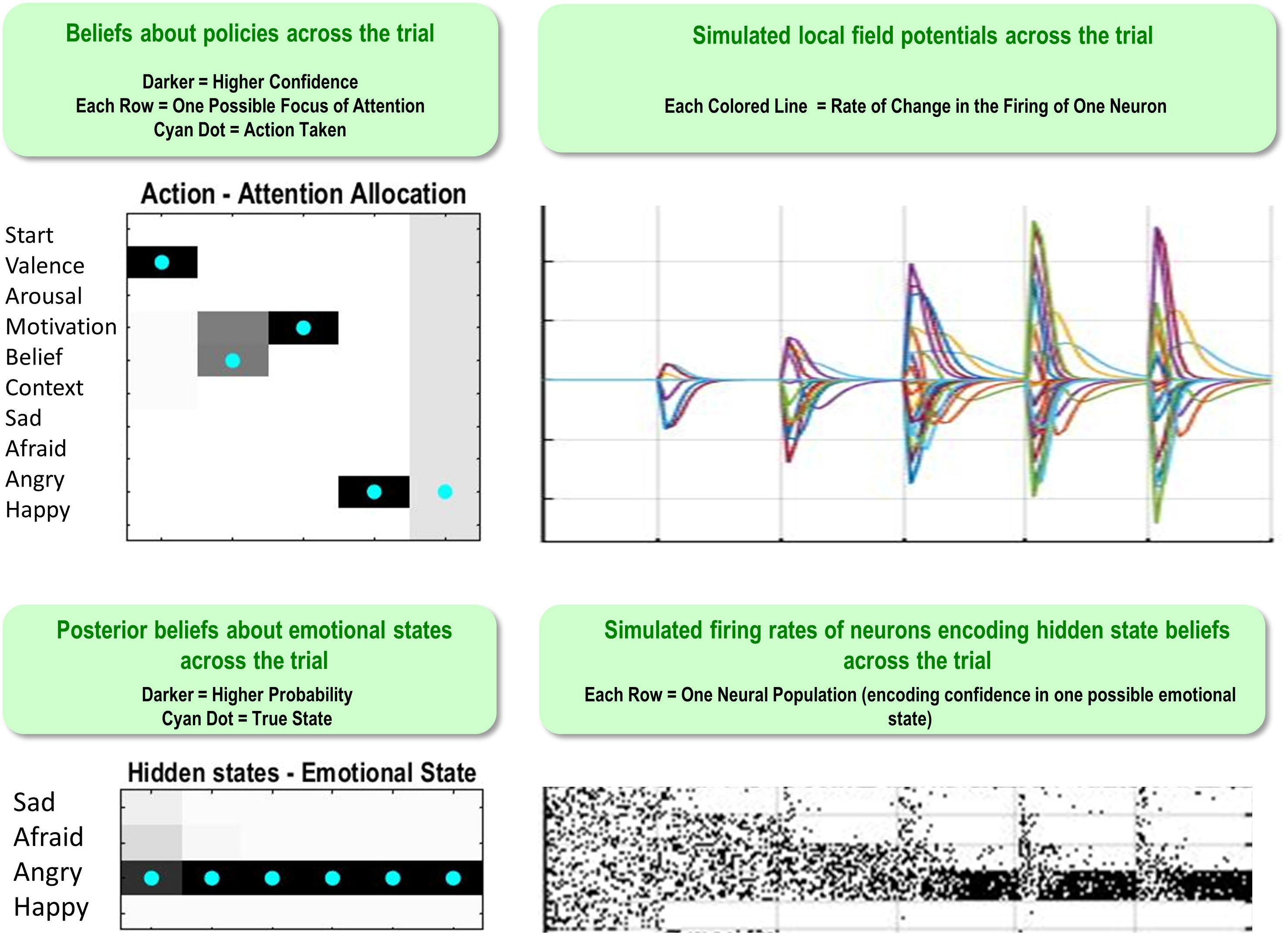
This figure illustrates a range of simulation results from one representative trial under high levels of A- and B-matrix precision (temperature parameter = 2 for each matrix). (Upper Left) Displays a sequence of chosen attentional shifts. The agent here chose a policy in which it attended to valence, then beliefs, then action motivation, and then chose to report that it was angry. Darker colors indicate higher confidence (probability estimates) in the model about its actions, whereas the cyan dot indicates the true action. (Bottom left) Displays the agent’s *posterior* beliefs about her emotional state across the trial. These posterior beliefs indicate that, at the end of the trial, the agent retrospectively inferred (correctly) that she was angry throughout the whole trial (i.e., despite not being aware of this until the fourth timestep, as indicated by the firing rates encoding confidence updates over time in the bottom right). (Lower Right/ Upper Right) Plots simulated neural responses in terms of single-neuron firing rates (raster plots) and local field potentials (rates of change in neural firing), respectively. Here each neuron’s activity codes the probability of occupying a particular hidden state at a particular point in time during the trial (based on the neural process theory depicted in figure 2; see (Friston, FitzGerald, et al., 2017)).

The simulations presented in Figures 4 and 5 make some cardinal points. First, it is fairly straightforward to simulate emotion processing in terms of emotional state inference. This rests upon a particular sort of generative model that can generate outcomes in multiple modalities. The recognition of an emotional state corresponds to the inversion of such models – and therefore necessarily entails multimodal integration. In other words, successfully disambiguating the most likely emotional state here requires consideration of the specific multimodal patterns of experience (i.e., incorporating interoceptive, exteroceptive, and proprioceptive sensations) that would be expected under each emotional state. We have also seen that this form of belief updating – or evidence accumulation – depends sensitively on what sort of evidence is actively attended. This equips the model of emotion concept representation with a form of mental action, which speaks to a tight link between emotion processing and attention to various sources of evidence from within the body – and beyond. Choices to shift attention vs. to self-report are respectively driven by the epistemic and pragmatic value of each allowable policy, such that pragmatic value gradually comes to drive the selection of reporting policies as the expected information gain of further attentional shifts decreases. The physiological plausibility of this emotion inference process has been briefly considered in terms of simulated responses. In the next section, we turn to a more specific construct validation, using empirical phenomenology from neurodevelopmental studies of emotion.

## Simulating the influence of early experience on emotional state inference and emotion concept learning

Having confirmed that our model could successfully infer emotional states – if equipped with emotion concepts – we are now in a position to examine emotion concept learning. Specifically, we investigated the conditions under which emotion concepts could be acquired successfully and the conditions under which this type of emotion learning and inference fails.

### Can emotion concepts be learned in childhood?

The first question we asked was whether our model could learn about emotions, if it started out with no prior beliefs about how emotions structure its experience. To answer this question, we first ran the model’s ‘A’ matrix (mapping emotion concepts to attended outcome information) through a softmax function with a temperature parameter of 0, creating a fully imprecise likelihood mapping. This means that each hidden emotional state predicted all outcomes equally (effectively, none of the hidden states within the emotion factor had any conceptual content). Then we generated 200 sets of observations (i.e., 50 for each emotion concept, evenly interleaved) based on the probabilistic state-outcome mappings encoded in the model described above (i.e., the “generative process”). That is, 50 interleaved learning trials for each emotion were generated by probabilistically sampling from a moderately precise version of the ‘A’ matrix distribution depicted in figure 3C (i.e., temperature parameter = 2). This resulted in 50 sets of observations consistent with the probabilistic mappings for each emotion (e.g., this entailed that roughly 50% of HAPPY trials involved observations of low vs. high arousal, whereas only roughly 1% of HAPPY trials involved the observation of social rejection, etc.). After the 200 learning trials, we then examined the changes in the model’s reporting accuracy over time. This meant that the agent, who began with no emotion knowledge (i.e., a fully uninformative ‘A’ matrix), observed patterns of observations consistent with each emotion (as specified above) at 50 timepoints spread out across the 200 trials and needed to learn these associations (i.e., learn the appropriate ‘A’ matrix mapping). This analysis was repeated at several levels of outcome (‘A’ matrix) and transition (‘B’ matrix) precision in the generative process – to explore how changes in the predictability or consistency of observed outcome patterns affected the model’s ability to learn. In this model, learning was implemented through updating (concentration) parameters for the model’s ‘A’ matrix after each trial. The model could also learn prior expectations for being in different emotional states, based on updating concentration parameters for its ‘D’ matrix after each trial (i.e., the emotional state it started in). For details of these free energy minimizing learning processes, please see (Friston et al., 2016).

We observed that the model could successfully reach 100% accuracy (with minor fluctuation) when both outcome and transition precisions in the generative process were moderately high (i.e., when the temperature parameters for the ‘A’ and ‘B’ matrix of the generative process were 2). The top panel in Figure 6 illustrates this by plotting the percentage accuracy across all emotions during learning over 200 trials (in bins of 10 trials), and for each emotion (in bins of 5). As can be seen, the model steadily approaches 100% accuracy across trials. The middle and lower panels of Figure 6 illustrate the analogous results when outcome precision and transition precision were lowered, respectively. The precision values chosen for these illustrations (‘A’ precision = 1.5, ‘B’ precision = 0.5) represent observed “tipping points” at which learning began to fail (i.e., at progressively lower precision values learning performance steadily approached 0% accuracy). As can be seen, lower precision in either the stability of emotions over time or the consistency between observations and emotional states confounded learning. Overall, these findings provide a proof of principle that this sort of model can learn emotion concepts, if provided with a representative and fairly consistent sample of experiences in its “childhood.”

**Figure 6.**
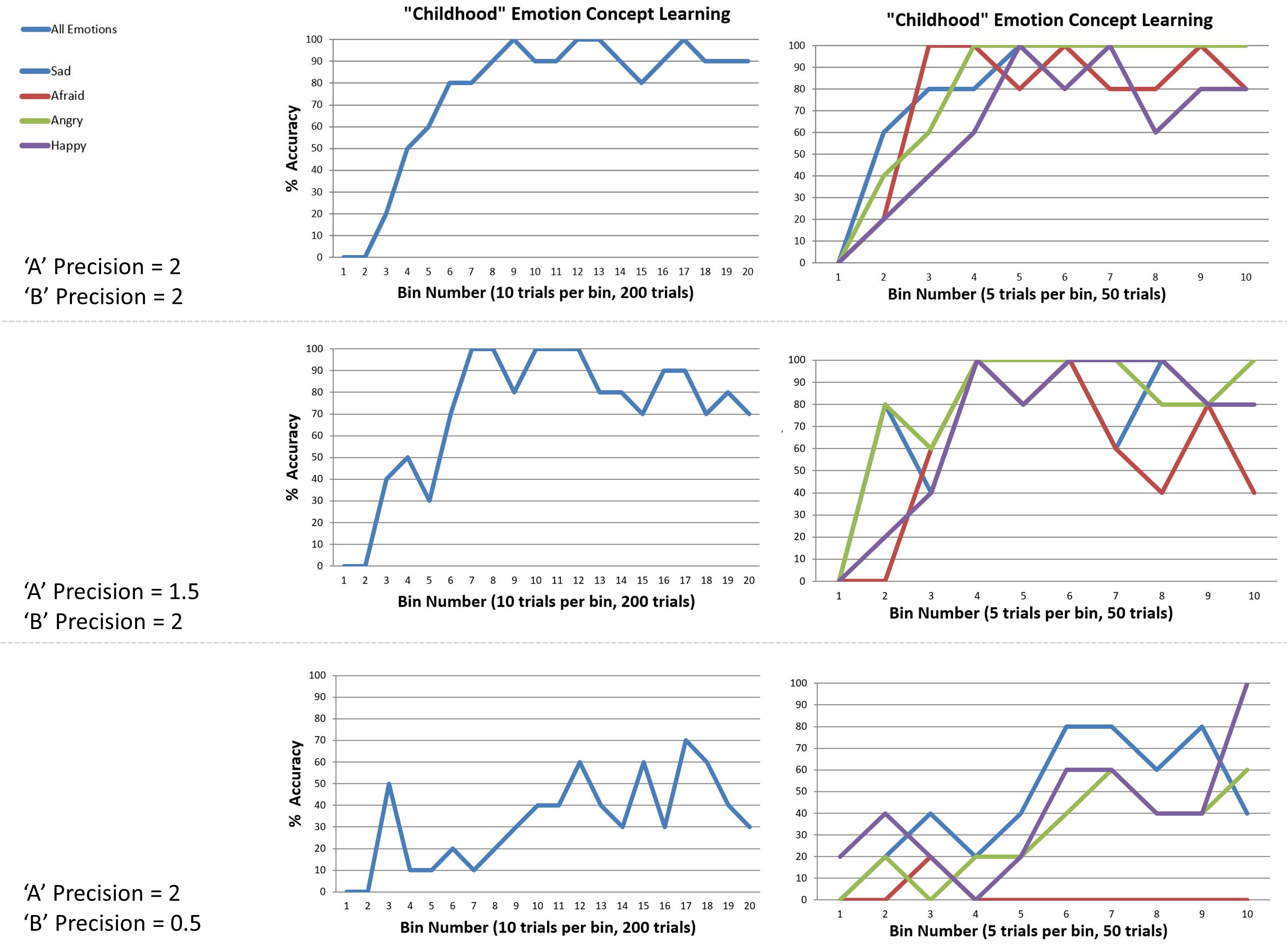
This figure illustrates simulated “childhood” emotion concept learning, in which the agent started out with no emotion concept knowledge (a uniform likelihood mapping from emotional states to outcomes) and needed to learn the correct likelihood mapping over 50 interleaved observations of the outcome patterns associated with each emotion concept (200 trials total). Left panels show changes in accuracy over time in 20-trial bins, at different levels of outcome pattern precision (i.e., ‘A’ and ‘B’ matrix precision, reflecting the consistency in the mapping between emotions and outcomes and the stability of emotional states over time, respectively). Right panels show the corresponding results for each emotion separately (5-trial bins). (Top) With moderately high precision (temperature parameter = 2 for both ‘A’ and ‘B’ matrices), learning was successful. (Middle and Bottom) With reduced ‘A’ precision or ‘B’ precision (respectively), learning began to fail.

### Can a new emotion concept be learned in adulthood?

We then asked whether a new emotion could be learned later, after others had already been acquired (e.g., as in adulthood). To answer this question, we again initialized the model with a fully imprecise ‘A’ matrix (temperature parameter = 0) and set the precision of the ‘A’ and ‘B’ matrices of the generative process to the levels at which “childhood” learning was successful (i.e., temperature parameter = 2 for each). We then exposed the model to 150 observations that only contained the outcome patterns associated three of the four emotions (50 for each emotion, evenly interleaved). We again allowed the model to accumulate experience in the form of concentration parameters for its ‘D’ matrix – allowing it to learn strong expectations for the emotional states it repeatedly inferred it was in. After these initial 150 trials, we then exposed the model to 200 further trials – using the outcome patterns under all four emotions (50 for each emotion, evenly interleaved). We then asked whether the emotion that was not initially necessary to explain outcomes could be acquired later, when circumstances change.

We first observed that, irrespective of which three emotions were initially presented, accuracy was high by the end of the initial 150 trials (i.e., between 80-100% accuracy for each of the 3 emotion concepts learned). The upper and middle panels of Figure 7 illustrate the accuracy over the subsequent 200 trials as the new emotion was learned. As can be seen in the upper left and right sections of Figure 7, ANGRY and SAD were both successfully learned. Interestingly, performance for the other emotions appeared to temporarily drop and then increase again as the new emotion concept was acquired (a type of temporary retroactive interference).

**Figure 7.**
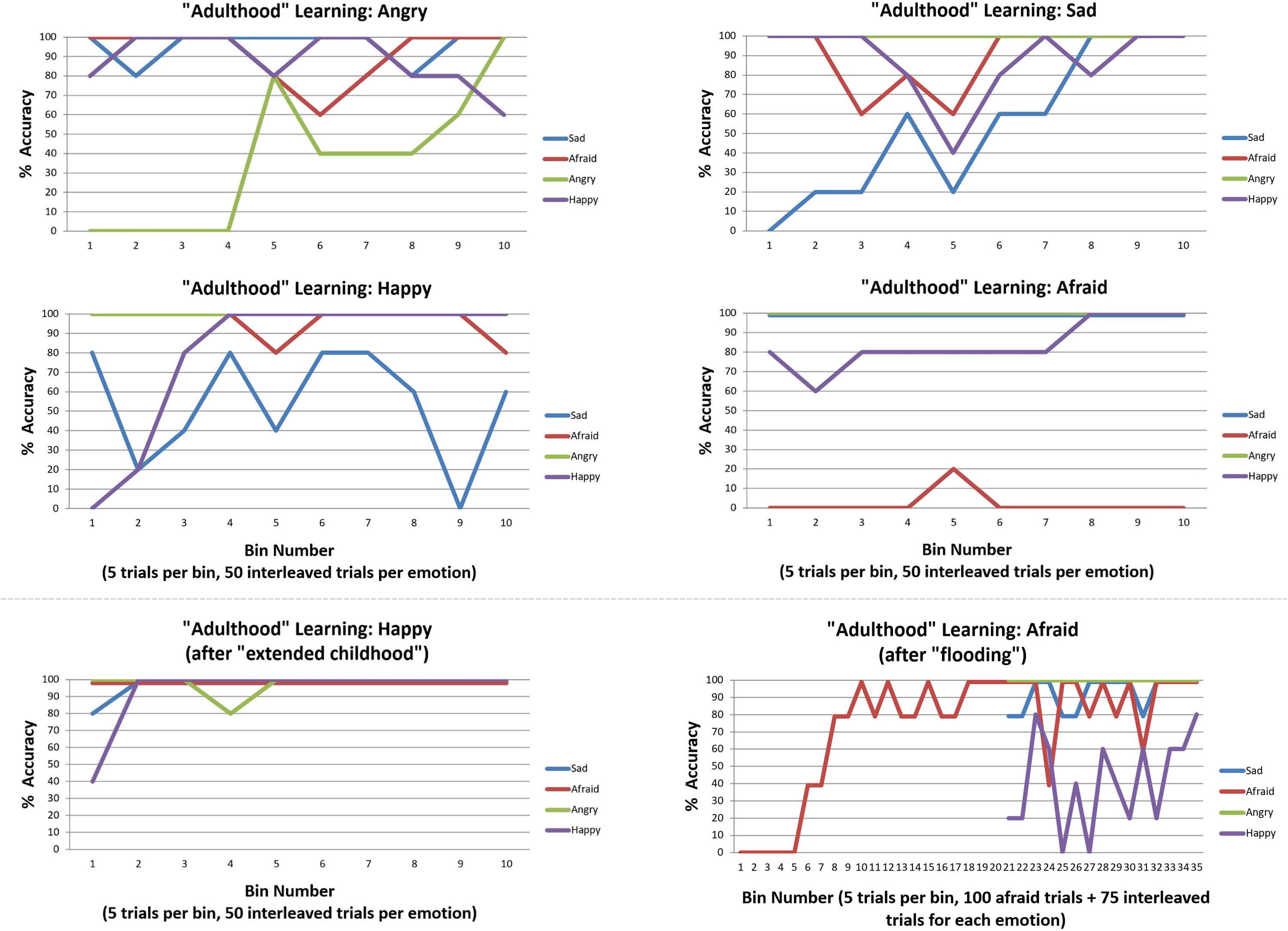
This figure illustrates emotion concept learning in “adulthood”, where three emotion concepts had already been learned (in 150 previous trials not illustrated) and a fourth was now needed to explain patterns of outcomes (over 200 trials, 50 of which involved the new emotion in an interleaved sequence). The top four plots illustrate learning for each of the four emotion concepts. ANGRY and SAD were successfully acquired. HAPPY was also successfully acquired, but interfered with previous SAD concept learning. As shown in the bottom left, this was ameliorated by providing the model with more “childhood” learning trials before the new emotion was introduced. AFRAID was not successfully acquired. This was (partially) ameliorated by first flooding the model with repeated observations of AFRAID-consistent outcomes before interleaving it with the other emotions, as displayed in the bottom right. All simulations were carried out with moderately high levels of precision (temperature parameter = 2 for both ‘A’ and ‘B’ matrices).

The middle left section of Figure 7 demonstrates that HAPPY could also be successfully learned; however, it appeared to interfere with prior learning for SAD. Upon further inspection, it appeared that SAD may not have been fully acquired in the first 150 trials (only reaching 80% accuracy near the end). We therefore chose to examine whether an “extended” or “emotionally enriched” childhood might prevent this interference, by increasing the initial learning trial number from 150 to 225 (75 interleaved exposures to SAD, ANGRY, and AFRAID outcome patterns). As can be seen in the lower left panel of Figure 7, HAPPY was quickly acquired in the subsequent 200 trials under these conditions, without interference with previous learning.

Somewhat surprisingly, the model was unable to acquire the AFRAID concept in its “adulthood” (Figure 7, middle right). To better understand this, for each trial we computed the expected evidence for each state, under the distribution of outcomes expected under the generative process (using bar notation to distinguish the process from the model) given a particular state, 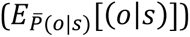. This was based on the reasoning that, if we treat the different emotional states as alternative models to explain the data, then the likelihood of data given states is equivalent to the evidence for a given state. Figure 8A plots the log transform of this expected evidence for each established emotion concept expected under the distribution of outcomes that would be generated if the ‘real’ emotional state were AFRAID (right panel) and contrasts it with the analogous plot for HAPPY (left panel), which was a more easily acquired concept (we took the logarithm of this expected evidence [or likelihood] to emphasize the lower evidence values; note that higher [less negative] values correspond to greater evidence in these plots).

As can be seen, in the case of HAPPY the three previously acquired emotion concepts had a relatively low ability to account for all observations, and so HAPPY was a useful construct in providing more accurate explanations of observed outcomes. In contrast, when learning AFRAID the model was already confident in its ability to explain its observations (i.e., the other concepts already had much higher evidence than in the case of HAPPY), and the “AFRAID” outcome pattern also provided moderate evidence for ANGRY and SAD (i.e., the outcome patterns between AFRAID and these other emotion concepts had considerable overlap). In figure 8B, we illustrate the ‘A’ and ‘D’ matrix values the agent had learned after the total 350 trials (i.e., 150 + 200, as described above), and an examplar trial in which the agent mistook fear for anger (which was the most common confusion). As can be seen, while the ‘A’ matrix mappings were learned fairly well, the agent had learned a strong prior expectation for ANGRY in comparison to its expectation for AFRAID. In the example trial the agent first attends to beliefs (observing other-blame) and then to the context (observing social rejection). These observations are consistent with both ANGRY and AFRAID; however, social rejection is more uniquely associated with ANGRY (i.e., AFRAID is also associated with crowded events, while ANGER is not). Combined with the higher prior expectation for ANGRY, the agent “jumps to conclusions” and becomes sufficiently confident to report ANGER (at which point she receives “incorrect” social feedback). Here, correct inference of AFRAID (i.e., disambiguating AFRAID from ANGRY) would have required that the agent also attend to her action tendencies (where she would have observed avoidance motivation) before deciding which emotion to report.

In this context, greater evidence for an unexplained outcome pattern would be required to “convince” the agent that its currently acquired concepts were not sufficient and that further information gathering (i.e., a greater number of attentional shifts) was necessary before becoming sufficiently confident to report her emotions. Based on this insight, we examined ways in which the model could be given stronger evidence that its current conceptual repertoire was insufficient to account for its observations. We first observed that we could improve model performance by “flooding” the model with an extended pattern of only AFRAID-consistent outcomes (i.e., 100 trials in a row), prior to reintroducing the other emotions in an interleaved fashion. As can be seen in Figure 7 (bottom right), this led to successful acquisition of AFRAID. However, it temporarily interfered with previous learning of the HAPPY concept. We also observed that by instead increasing the number of AFRAID learning trials from 200 to 600, the model eventually increased its accuracy to between 40% and 80% across the last 10 bins (last 50 trials) – indicating that learning could occur, but at a much slower rate.

**Figure 8A.**
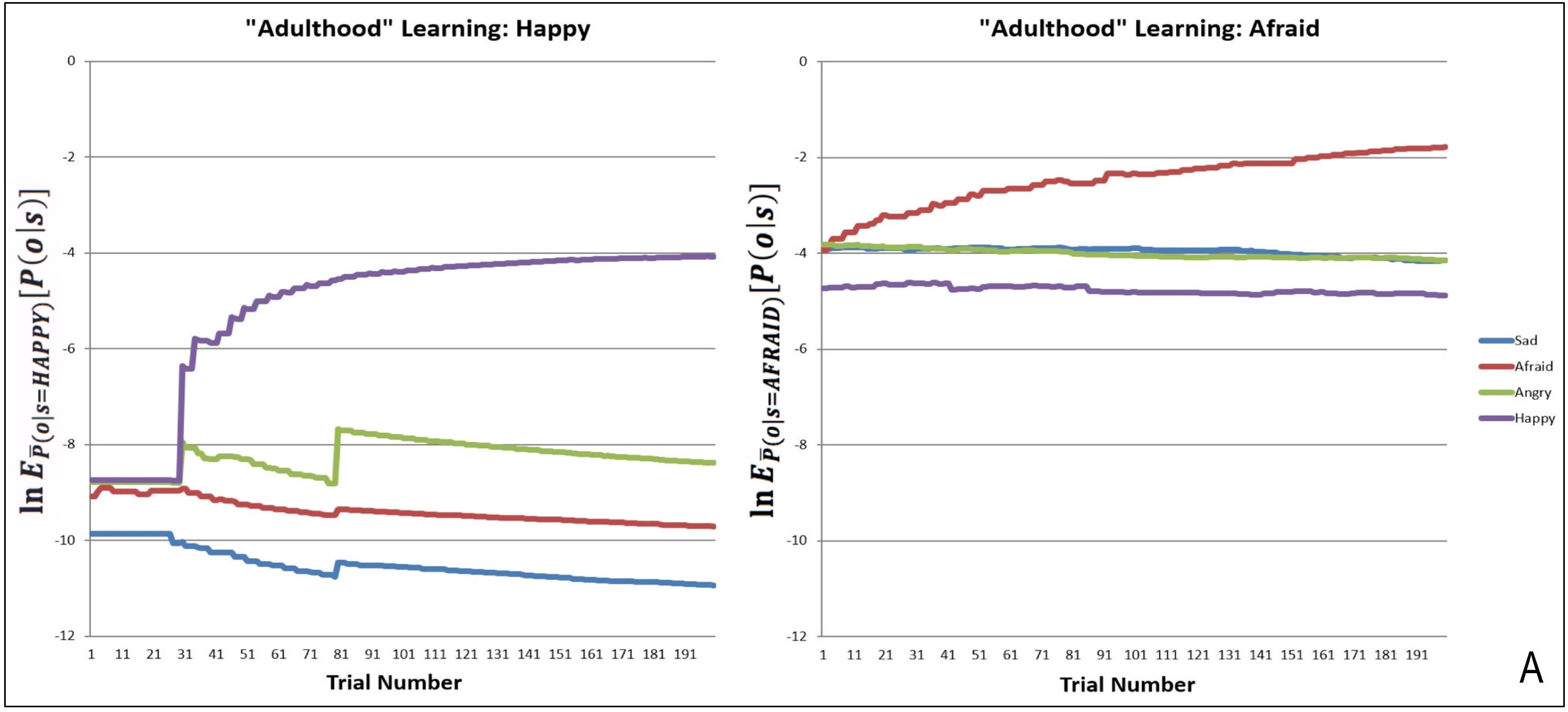
(Left) The log expected probability of outcomes under the HAPPY outcome distribution, given each of the four emotion concepts in the agent’s model (i.e., given the ‘A’ matrix it had learned) at each of 200 learning trials, *E_P̄(o|s = HAPPY)_ [P(o|s)]* (note: the bar over P indicates the generative process distribution). (Right) The analogous results for the AFRAID outcome pattern. These plots illustrate that HAPPY may have been more easily acquired than AFRAID because the agent was less confident in the explanatory power of its current conceptual repertoire when it began to observe the HAPPY outcome pattern than when it began to observe the AFRAID outcome pattern. This also shows that the AFRAID outcome pattern provided some evidence for SAD and ANGRY in the agent’s model, likely due to outcome pattern overlap between AFRAID and these other two concepts.

**Figure 8B.**
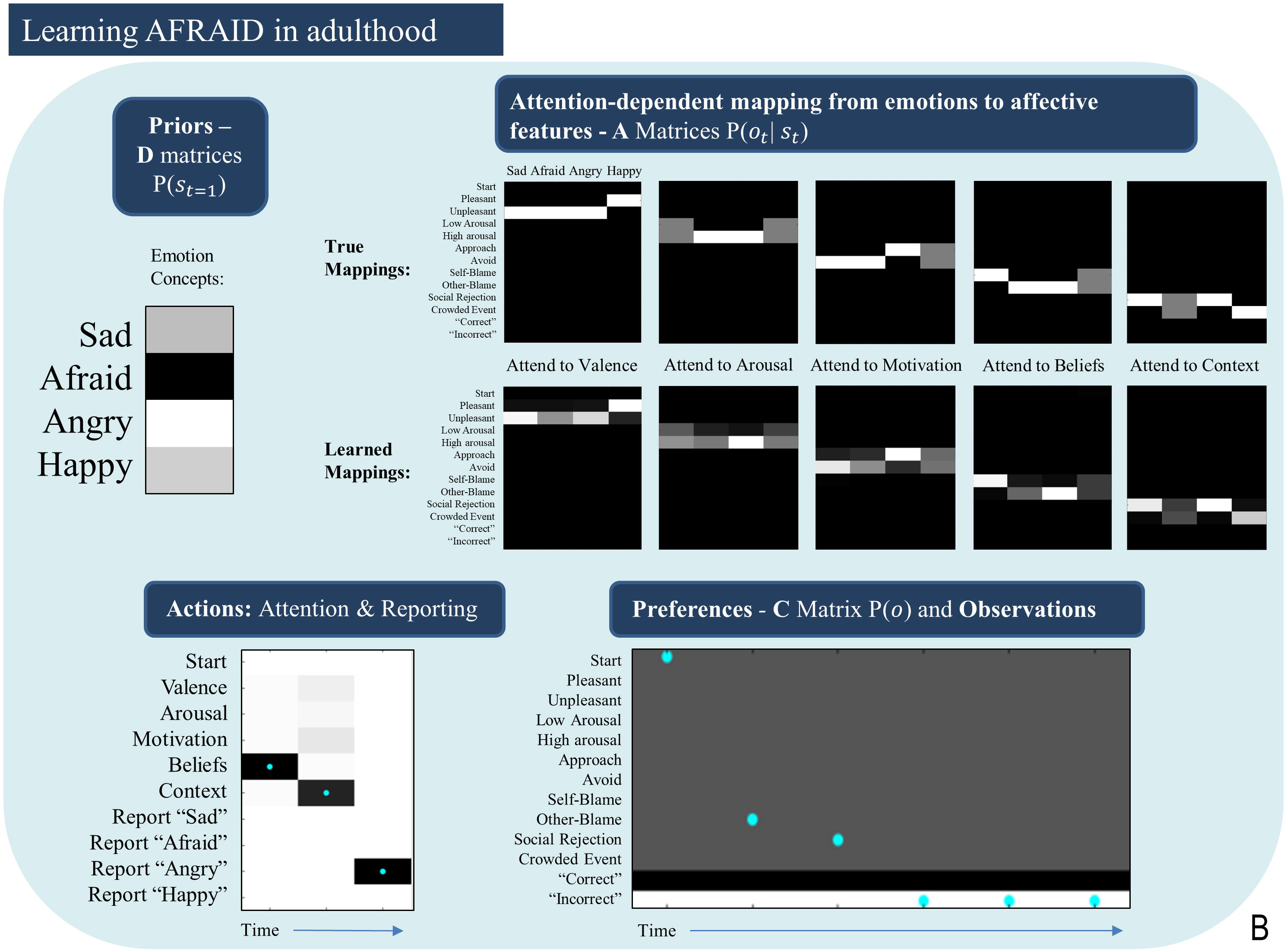
Illustrated the priors (D) and likelihoods (A) learned after 50 observations of SAD, ANGRY, and HAPPY (150 trials total) followed by 200 trials in which SAD, AFRAID, ANGRY, and HAPPY were each presented 50 times. As can be seen, while the ‘A’ matrices are fairly well learned, the agent acquired a strong prior expectation for ANGRY. As shown in the example trial in the bottom left, this led the agent to “jump to conclusions” and report ANGER on AFRAID trials after making 2 observations consistent with both ANGRY and AFRAID (other-blame and social rejection, as shown in the bottom right). In this case, the agent would have needed to attend to her motivated actions (avoid) to correctly infer AFRAID.

Overall, these results confirmed that a new emotion concept could be learned in synthetic “adulthood,” as may occur, for example, in psycho-educational interventions during psychotherapy. However, these results also demonstrate that this type of learning can be more difficult. These results therefore suggest a kind of “sensitive period” early in life where emotion concepts are more easily acquired.

### Can maladaptive early experiences bias emotion conceptualization?

The final question we asked was whether unfortunate early experiences could hinder our agent’s ability to adaptively infer and/or learn about emotions. Based on the three-process model (R. Smith, Killgore, et al., 2018), it has previously been suggested that at least two mechanisms could bring this about:

1. Impoverished early experiences (i.e., not being exposed to the different patterns of observations that would facilitate emotion concept learning).
2. Having early experiences that reinforce maladaptive cognitive habits (e.g., selective attention biases), which can hinder adaptive inference (if the concepts have been acquired) and learning (if the concepts have not yet been acquired).

We chose to examine both of these possibilities below.

#### Non-representative early emotional experiences

To examine the first mechanism (involving the maladaptive influence of “unrepresentative” early experiences), we used the same learning procedure and parameters described in the previous sections. In this case, however, we exposed the agent to 200 outcomes generated by a generative process where one emotion was experienced 50 times more often than others. Specifically, we examined the cases of a childhood filled with either chronic fear/threat or chronic sadness, as a potential means of simulating the effects of continual childhood abuse or neglect (the sadness simulations might also be relevant to chronic depression over several years). We then examined the model’s ability to learn to infer new emotions in a subsequent 200 trials.

In general, we observed that primarily experiencing fear or sadness during childhood (which could also be thought of as undifferentiated in the sense that they could not be contrasted with other emotions) led the agent to have notable difficulties in learning new emotions later in life. These results were variable upon repeated simulations with different emotions (e.g., verbal reporting continually fluctuated between high and low levels of accuracy for some emotions, while accuracy remained near 0% for others, while yet others were well acquired). For example, in one representative simulation, in which the agent primarily experienced fear during childhood, reporting accuracy continually varied for HAPPY (45% accuracy in the final 20 trials), remained at 0% for ANGRY, remained at 100% for AFRAID, and was stable at 100% for SAD). Whereas primarily experiencing sadness in childhood during a representative simulation led to 0% accuracy for HAPPY, continually varying accuracy for ANGRY and AFRAID (55% accuracy in the final 20 trials for each), and stable high accuracy for SAD (95% accuracy in final 20 trials). Similar patterns of (highly variable) results were observed when performing the same simulations with the other two emotions.

Unlike the results shown in figure 8B – in which likelihood mappings were fairly well acquired (and precise prior expectations for specific emotions hindered correct inference) – poor performance was here explained primarily by poorly acquired likelihood mappings (i.e., the content of the other emotions concepts was often not learned). Figure 9 illustrates this by presenting the ‘A’ matrices learned by the agent after childhoods dominated by either fear or sadness. As can be seen there, the likelihood mappings do not strongly resemble the true mapping within the generative process. These results in general support the notion that having unrepresentative or insufficiently diverse early emotional experiences could hinder later learning.

**Figure 9.**
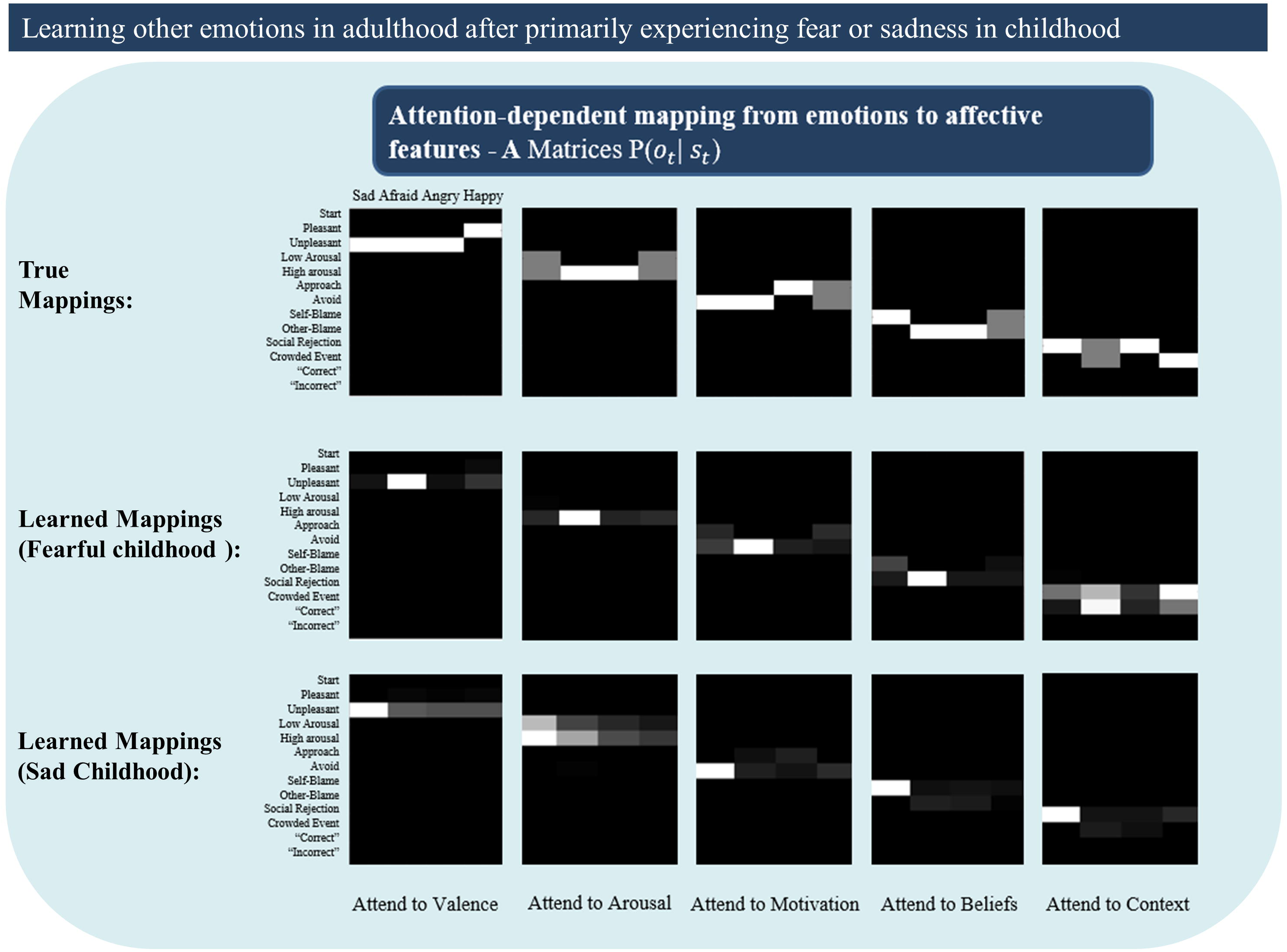
Illustrates poorly learned emotion concepts (likelihood mappings or ‘A’ matrices) in adulthood due to a childhood primarily characterized by either fear or sadness. This could be thought of as simulating early adversity involving continual abuse or neglect, or perhaps cases of chronic depression. See text for more details.

#### Maladaptive Attention Biases

To examine the second mechanism proposed by the three-process model (involving maladaptive patterns in habitual attention allocation), we equipped the model’s ‘E’ matrix with high prior expectations over specific policies, which meant that it was 50 times more likely to attend to some information and not to other information. This included: (i) an “external attention bias,” where the agent had a strong habit of focusing on external stimuli (context) and its beliefs about self- and other-blame; (ii) an “internal attention bias,” where the agent had a strong habit of only attending to valence and arousal; and (iii) a “somatic attention bias,” where the agent had a strong habit to attend only to its arousal level and the approach vs. avoid modality.

Figure 10 shows how these different attentional biases promote false inference. On the left, the true state is AFRAID, and the externally focused agent first attends to the stimulus/context (social rejection) and then to its beliefs (other-blame); however, without paying attention to its motivated action (avoid), it falsely reports that it feels ANGRY instead of AFRAID (note that, following feedback, there is a retrospective inference that afraid was more probable; similar retrospective inferences after feedback are also shown in the other two examples in Figure 10). In the middle, the true state is ANGRY, and the internally focused agent first attends to valence (unpleasant) and then to arousal (high); however, without paying attention to its action tendency (approach), it falsely reports that it is AFRAID. On the right, the true state is SAD, and the somatically focused agent attends to its motivated action (avoid) and to its arousal (high); however, without attending to its beliefs (self-blame) it falsely reports that it is AFRAID instead of SAD.

Importantly, these false reports occur in an agent that has already acquired very precise emotion concepts. Thus, this does not represent a failure to learn, but simply the effect of having learned poor habits for mental action.

**Figure 10.**
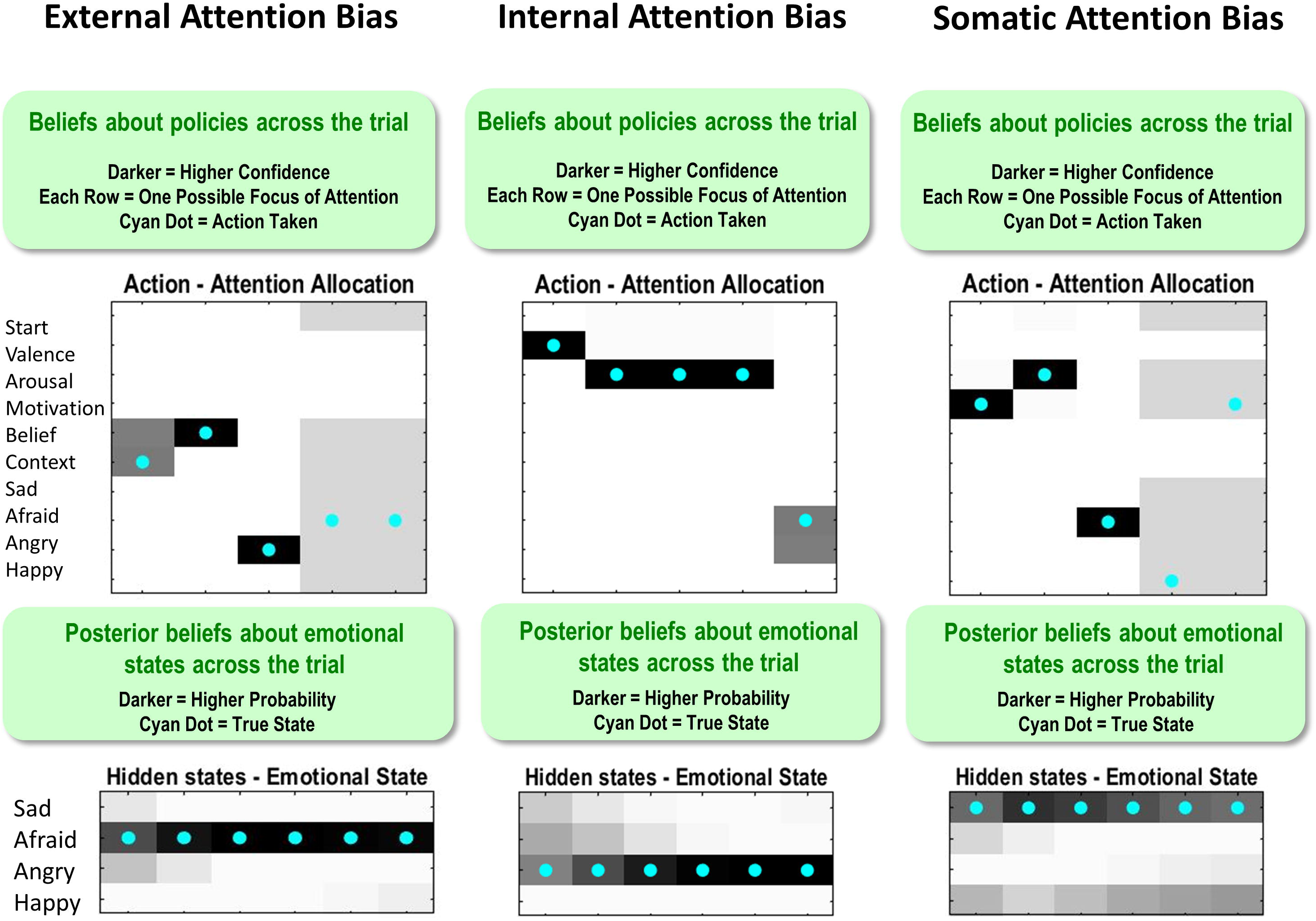
This figure illustrates single trial effects of different attentional biases, each promoting false inference in a model that has already acquired precise emotion concepts (i.e., temperature parameter = 2 for both ‘A’ and ‘B’ matrices). External attention bias was implemented by giving the agent high prior expectations (‘E’ matrix values) that she would attend to the context and to her beliefs. Internal attention bias was implemented by giving the agent high prior expectations that she would attend to valence and arousal. Somatic attention bias was implemented by giving the agent high prior expectations that she would attend to arousal and action motivation. See the main text for a detailed description of the false inferences displayed for each bias. Note: the bottom panels indicate *posterior* beliefs, illustrating the agent’s *retrospective* beliefs at the end of the trial about her emotional state throughout the trial. Also note that the agent’s actions in the upper left and upper right after reporting should be ignored (i.e., they are simply an artifact of the trial continuing after the agent’s report, when there is no goal and therefore no confidence in what to do).

The results of these examples were confirmed in simulations of 40 interleaved emotion trials (10 per emotion) in an agent who had already acquired precise emotion concepts (temperature parameter = 2 for both ‘A’ and ‘B’ matrices; no learning). In these simulations, we observed that the externally focused agent had 100% accuracy for SAD, 10% accuracy for AFRAID, 100% accuracy for ANGRY, and 60% accuracy for HAPPY. The internally focused agent had 100% accuracy for SAD, 50% accuracy for AFRAID, 60% accuracy for ANGRY, and 100% accuracy for HAPPY. The somatically focused agent had 10% accuracy for SAD, 100% accuracy for AFRAID, 100% accuracy for ANGRY, and 0% accuracy for HAPPY. Thus, without adaptive attentional habits, the agent was prone to misrepresent its emotions.

In our final simulations, we examined how learning these kinds of attentional biases in childhood could hinder emotion concept learning. To do so, we used the same learning procedure describe in section 3.1. However, in this case we simply equipped the agent with the three different attentional biases (‘E’ matrix prior distributions over policies) and assessed its ability to learn emotion concepts over the 200 trials. The results of these simulations are provided in Figure 11. A somatic attention bias primarily allowed the agent to learn two emotion concepts, which corresponded to ANGER and FEAR. However, it is worth highlighting that the low accuracy for the other emotions means that their respective patterns were subsumed under the first two. Thus, it is more accurate to say that the agent learned two affective concepts, which largely predicted only approach vs. avoidance. In contrast, the internally biased agent easily acquired the distinction between pleasant (HAPPY) and unpleasant emotions (all lumped into ANGER) and began to learn a third concept that distinguished between low vs. high arousal (i.e., SAD vs. ANGRY). However, it did not conceptualize the distinction between approach and avoidance (i.e., ANGRY vs. AFRAID). Lastly, the externally focused agent was somewhat labile in its concept acquisition; by the end, it could not predict approach vs. avoidance, but it possessed externally focused concepts with content along the lines of “the state I am in when I’m socially rejected and think it’s my fault vs. someone else’s fault” (SAD vs. ANGRY) and “the state I am in when at a crowded event” (HAPPY). These results provide strong support for the role of attentional biases in subverting emotional awareness.

**Figure 11.**
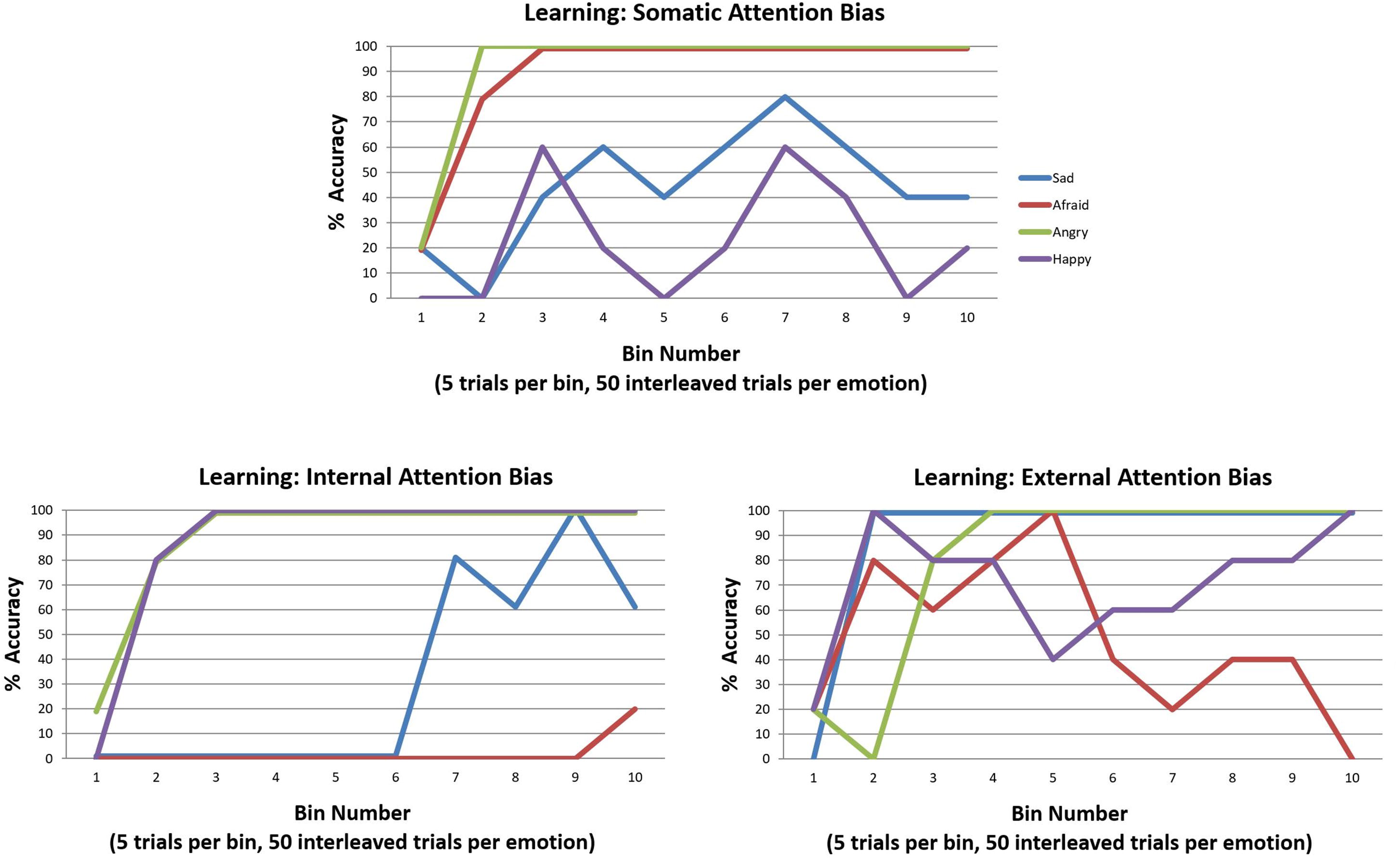
Illustration of the simulated effects of each of the three attention biases (described in the legend for Figure 10) on emotion concept learning over 200 trials (initialized with a model that possessed a completely flat ‘A’ matrix likelihood distribution). Each bias induces aberrant learning in a different way, often leading to the acquisition of two more coarse-grained emotion concepts as opposed to the four fine-grained concepts distinguished in the generative process. See the main text for a detailed description and interpretation.

## Discussion

The active inference formulation of emotional processing we have presented represents a first step toward the goal of building quantitative computational models of the ability to learn, recognize, and understand (be “aware” of) one’s own emotions. Although this is clearly a toy model, it does appear to offer some insights, conceptual advances, and possible predictions.

First, in simulating differences in the precision (specificity) of emotion concepts, some intuitive but interesting phenomena emerged. As would be expected, differences in the specificity of the content of emotion concepts – here captured by the precision of the likelihood mapping from states to outcomes (i.e., the precision of what pattern of outcomes each emotion concept predicted) – led to differences in inferential accuracy. This suggests that, as would be expected, those with more precise emotion concepts would show greater understanding of their own affective responses. Perhaps less intuitively, beliefs about the stability of emotion concepts – here captured by the precision of expected state transitions – also influenced inferential accuracy. This predicts that a belief that emotional states are more stable (less labile) over time would also facilitate one’s ability to correctly infer what they are feeling. This appears consistent with the low levels of emotional awareness or granularity observed in borderline personality disorder, a disorder characterized by emotional instability (Levine et al., 1997; Suvak et al., 2011).

Next, in simulating emotion concept learning, a few interesting insights emerged. Our simulations first confirmed that emotion concepts could successfully be learned, even when their content was cast (as done here) as complex, probabilistic, and highly overlapping response patterns across interoceptive, proprioceptive, exteroceptive, and cognitive domains. This was true when all emotion concepts needed to be learned simultaneously (as in childhood; see (Hietanen, Glerean, Hari, & Nummenmaa, 2016; Widen & Russell, 2008)), and was also true when a single new emotion concept was learned after others had already been acquired (as in adulthood during psycho-educational therapeutic interventions; e.g., see (Barlow et al., 2016; Burger et al., 2016; Hayes & Smith, 2005; Lumley et al., 2017)).

These results depended on whether the observed outcomes during learning were sufficiently precise and consistent. One finding worth highlighting was that emotion concept learning was hindered when the precision of transitions among emotional states was too low. This result may be relevant to previous empirical results in populations known to show reduced understanding of emotional states, such as those with autism (Erbas, Ceulemans, Boonen, Noens, & Kuppens, 2013; Silani et al., 2008) and those who grow up in socially impoverished or otherwise adverse (unpredictable) environments (Colvert et al., 2008; Lane et al., 2018). In autism, it has been suggested that overly imprecise beliefs about state transitions may hinder mental state learning, because such states require tracking abstract behavioral patterns over long timescales (Haker, Schneebeli, & Stephan, 2016; Lawson, Mathys, & Rees, 2017; Lawson, Rees, & Friston, 2014). Children who grow up in impoverished environments may not have the opportunity to interact with others to observe stable patterns in other’s affective responses; or receive feedback about their own (Lane et al., 2018; Pears & Fisher, 2005; R. Smith, Killgore, et al., 2018). Our results successfully reproduce these phenomena – which represent important examples of mental state learning that may depend on consistently observed outcome patterns that are relatively stable over time.

As emotion concepts are known to differ in different cultures (Russell, 1991), our model and results may also relate to the learning mechanisms allowing for this type of culture-specific emotion categorization learning. Specifically, the “correct” and “incorrect” social feedback in our model could be understood as linguistic feedback from others in one’s culture (e.g., a parent labeling emotional reactions for a child using culture-specific categories). If this feedback is sufficiently precise, then emotion concept learning could proceed effectively – even if the probabilistic mapping from emotion categories to other perceptual outcomes is fairly imprecise (i.e., which appears to be the case empirically; (Barrett, 2006, 2017)).

Another insight worth highlighting was that learning was more difficult when the agent had already acquired previous concepts but entertained the possible existence of new emotions it had not already learned. An interesting observation was that, in some cases (e.g., learning SAD), new learning temporarily interfered with old learning before being fully integrated into the agent’s cognitive repertoire (an effect – termed retroactive interference – that has been extensively studied empirically within learning and memory research; (Darby & Sloutsky, 2015; Martínez, Villar, Ballarini, & Viola, 2014)). In the case of learning HAPPY, we found that more extensive learning in the model’s “childhood” was necessary to prevent this type of interference, with respect to previous acquisition of the concept SAD. A second interesting observation was that AFRAID was very difficult to learn after the other concepts were fully acquired. This appeared to be because the agent had already learned to be highly confident about the explanatory power of her current conceptual repertoire (combined with the fact that AFRAID had considerable outcome overlap with SAD and ANGRY). It was necessary to provide the model with a persistent “flooding” of observations consistent with the new emotion to reduce its confidence sufficiently to acquire a new concept. It is not clear that this type of flooding is realistic, but perhaps resembles the extended periods of fear that occur during exposure-based behavioral therapies (Cooper, Clifton, & Feeny, 2017). We should also emphasize that, due to the oversimplified nature of the mappings between emotion concepts and affective response features in our simulations, the difficulties observed in learning these specific emotions should not be taken too seriously. However, these overall results do predict that emotion concept learning in general should be more difficult in adulthood, and that emotion learning may have a kind of “sensitive period” in childhood (as supported by previous empirical findings; e.g., (Colvert et al., 2008; Pears & Fisher, 2005)).

The manner in which concept learning was implemented in these simulations may also have more general implications (for considerations of this approach to concept learning more generally, see (R. Smith, Schwartenbeck, Parr, & Friston, 2019)). Typically in Active Inference simulations, the state space structure of a model is specified in advance (e.g., (Mirza et al., 2016; Parr & Friston, 2017b; Schwartenbeck, FitzGerald, Mathys, Dolan, & Friston, 2015)). Our model was instead equipped with “blank” hidden states devoid of content (i.e., these states started out predicting all outcomes with roughly equal probability in the simulated learning). Over multiple exposures to the observed outcomes, these blank hidden states came to acquire conceptual content that captured distinct statistical patterns in the lower-level affective response components of the model. In some current neural process theories (Bastos et al., 2012; Friston, FitzGerald, et al., 2017; Friston, Rosch, Parr, Price, & Bowman, 2018; Parr & Friston, 2018), distinct cortical columns are suggested to represent distinct hidden states. Under such theories, our learning model would suggest that the brain might contain “reserve” cortical columns available to capture new patterns of lower-level covariance if/when they begin to be observed in interaction with the world. To our knowledge, no direct evidence of such “reserve neurons” has been observed, although the generation of new neurons (with new synaptic connections) is known to occur in the hippocampus (Chancey et al., 2013). There is also the well-known phenomenon of “silent synapses” in the brain, which can persist into adulthood and become activated when new learning becomes necessary (e.g., see (Chancey et al., 2013; Funahashi, Maruyama, Yoshimura, & Komatsu, 2013; Kerchner & Nicoll, 2008)). Another interesting consideration is that, during sleep, it appears that many (but not all) synaptic strength increases acquired in the previous day are attenuated (Tononi & Cirelli, 2014). This has been suggested to correspond to a process of Bayesian model reduction, in which redundant model parameters are identified and removed to prevent model over-fitting and promote selection of the most parsimonious model that can successfully account for previous observations (Hobson & Friston, 2012). This also suggests that increases in “reserve” representational resources available for state space expansion (as in concept learning) could perhaps occur after sleep. In short, the acquisition of new concepts, emotion-related or otherwise, speaks to important issues in structure learning. The approach used here offers one solution to the question of how to expand a model, which could complement work on strategies for reducing a model (Friston, Lin, et al., 2017).

Although the neural process theory associated with active inference is cast at the level of canonical microcircuits and message passing, and therefore does not make a priori predictions about the brain regions that implement the emotion-related processes in our model, it nonetheless can afford empirical testing of macro-anatomical correlates. That is, this process theory can be used to generate predicted neural response time courses during emotional state inference and emotion concept learning in our model, and the macro-anatomical correlates of these time courses can then be established using neuroimaging methods. At present, the three-process model (R. Smith, Killgore, et al., 2018), and supporting evidence (McRae et al., 2008; R. Smith, Alkozei, et al., 2017; R. Smith et al., 2015; R. Smith, Kaszniak, et al., 2019; R. Smith, Lane, Alkozei, et al., 2018; R. Smith, Lane, et al., 2017; R. Smith, Lane, Sanova, et al., 2018; R. Smith, Quinlan, et al., 2019; R. Smith, Sanova, et al., 2018), has identified a number of large-scale networks that plausibly implement the processes we have simulated – and could therefore provide a priori hypotheses for future studies along these lines (Barrett & Satpute, 2013; Yeo et al., 2011). For example, “limbic network” regions (including orbitofrontal cortex and amygdala, among others), “salience network” regions (including the anterior insula and dorsal anterior cingulate, among other) and somatomotor/posterior insula regions all appear to be involved in generating affective responses and representing either visceral, somatic, or proprioceptive states at a perceptual level. Regions of the paralimbic cortex (e.g., “default mode network,” with major hubs in the medial prefrontal cortex and posterior cingulate) are in turn most strongly implicated in conceptual inference (Binder, Desai, Graves, & Conant, 2009) – such as the emotion concept representation processes simulated here. Thus, activity in a number of distinct brain regions/networks would be expected to show associations with distinct belief updating processes in our model.

A final insight offered by our model pertains to the possibility that maladaptive emotional state inference could be due to early experience. We demonstrated this in two ways. First, we simulated exposure to a large number of single emotion-provoking situations in childhood, promoting precise and highly engrained prior expectations for being in a single emotional state, as well as preferential learning of the respective outcome patterns for that state over others. We found that different kinds of “unrepresentative” (single-emotion) outcome patterns in early experience (e.g., chronic fear or sadness in childhood abuse/neglect or sever chronic depression) prevented learning other emotion concepts in somewhat inconsistent ways in repeated simulations. Overall, however, these results supported the idea that later emotional state inferences and emotion concept learning could be compromised by this type of maladaptive early experience. This could potentially relate to cognitive bias learning, such as the negative interpretation biases characteristic of mood and anxiety disorders (which have been interpreted within computational frameworks; (Mogg & Bradley, 2005; R. Smith, Akozei, Killgore, & Lane, 2017)).

Second, we examined the possibility that maladaptive cognitive habits could hinder emotional awareness. Here, we demonstrated that such habits can promote false emotional state inference and can hinder emotion concept learning. Specifically, we found that different types of external, internal, and somatic biases led to the acquisition of coarser-grained emotion concepts that failed to distinguish between various elements of affective responses. Aside from its relevance to cognitive biases more generally, these results could also explain certain empirical phenomena in emotional awareness research, such as the finding that males tend to score lower on emotional awareness measures than females (Wright et al., 2017). Specifically, while a genetic contribution to such findings is possible, it is also known that many cultures reinforce emotion avoidance in boys more than in girls in childhood (Chaplin, Cole, & Zahn-Waxler, 2005; Diener & Lucas, 2004; Fivush, Brotman, Buckner, & Goodman, 2000), and can promote beliefs that paying attention to emotions is a sign of weakness or that emotional information simply carries little pragmatic value. This type of learning could plausibly reinforce biased patterns of attention similar to those simulated here. Thus, our simulations suggest an interesting, testable mechanism by which such (potentially socialization-based) differences may arise.

In closing, it is important to note that this model is deliberately simple and is meant only to represent a proof of principle that emotion inference and learning can be modeled within a neurocomputational framework from first principles. We chose a particular pattern of state-outcome mappings to simulate the content of emotion concepts, but this is unlikely to represent a fully accurate depiction of human emotion concepts or the outcomes they predict. Human emotion concepts likely draw on much higher-dimensional patterns of somatic and visceral sensations, behavioral motivations, and cognitive appraisal patterns. There are also “secondary” emotion concepts like jealousy or embarrassment, which may require including more specific context and appraisal observations in a model (e.g., observing a lover with a competing suitor, observing oneself committing actions that break social norms, etc.). Further, human agents (or at least some of them) are likely to have a much richer space of both emotion and non-emotion concepts available for explaining their patterns of internal experience in conjunction with other beliefs and exteroceptive evidence (e.g., a pattern of low arousal, unpleasant valence, and avoidance in many contexts could also be explained by the concept of sickness rather than sadness; see (R. Smith, Lane, et al., 2019)). A more complete model would take into account many different possible conceptual interpretations of this sort. In addition, our simulations only attempted to capture the second process within the three-process model (i.e., affective response representation; (R. Smith, Killgore, et al., 2018)). Incorporating the other two processes (affective response generation and conscious access) would undoubtedly induce additional dynamics (including explicit brain-body interactions) that could alter or nuance the simulation results we have provided. Modeling these additional processes will be an important goal of future work (see (R. Smith, Lane, et al., 2019)).

A final more general limitation with this type of modeling is that, in its current form, there are limited means of evaluating how well it represents the true form of emotional state inference and emotion concept learning implemented in the human brain. Here, we have focused on reproducing and validating a minimal model that evinces emotional state inference and learning within the active inference framework. Crucially, this model has – by construction – a construct validity with the three-process model and associated empirical evidence. As noted above, external validation of the model’s ability to capture human brain processes will be an important next step, and can be done, for example, by examining whether the simulated neural responses we have presented are observable within particular brain regions during future neuroimaging studies of attending to – and reporting – one’s own emotions (e.g., (Gusnard, Akbudak, Shulman, & Raichle, 2001; Lane, Fink, Chua, & Dolan, 1997; R. Smith, Fass, & Lane, 2014; R. Smith, Lane, Alkozei, et al., 2018; R. Smith, Lane, Sanova, et al., 2018)).

With these limitations in mind, however, this approach to computationally modeling emotion-related processes appears promising with respect to the initial insights it can offer. It can illustrate selective information integration in the service of conceptual inference, it can successfully simulate concept learning and some of its known vulnerabilities, and it can highlight maladaptive interactions between cognitive habits, early experience, and the ability to understand and be aware of one’s own emotions later in life, all of which may play important roles in the development of emotional pathology. Finally, it highlights the potential for future empirical work in which tasks could be adapted to the broad structure of such models, which would allow investigation of individual differences in emotion processing as well as its neural basis. In other words, once we have a validated model of these emotion-related processes – at the subjective and neuronal level – we can, in principle, fit the model to observed responses and thereby phenotype subjects in terms of their emotion-related beliefs states (Schwartenbeck & Friston, 2016).

### Software note

Although the generative model – specified by the various matrices described in this paper – changes from application to application, the belief updates are generic and can be implemented using standard routines (here **spm_MDP_VB_X.m**). These routines are available as Matlab code in the latest version of SPM academic software: http://www.fil.ion.ucl.ac.uk/spm/. The simulations in this paper can be reproduced (and customized) via running the Matlab code included here in the supplementary material (**Emotion_learning_model.m**).

## Supporting information

Emotion_learning_model.m

## References

Bagby, R., Parker, J., & Taylor, G. (1994a). The twenty-item Toronto Alexithymia Scale—I. Item selection and cross-validation of the factor structure. Journal of Psychosomatic Research, 38(1), 23–32.

Bagby, R., Parker, J., & Taylor, G. (1994b). The twenty-item Toronto Alexithymia Scale—II. Convergent, discriminant, and concurrent validity. Journal of Psychosomatic Research, 38(1), 33–40.

Barchard, K., & Hakstian, A. (2004). The nature and measurement of emotional intelligence abilities; basic dimensions and their relationships with other cognitive abilities and personality variables. Educational and Psychological Measurement, 64, 437–462.

Barlow, D., Allen, L., & Choate, M. (2016). Toward a Unified Treatment for Emotional Disorders - Republished Article. Behavior Therapy, 47(6), 838–853. https://doi.org/10.1016/j.beth.2016.11.005

Barrett, L. (2006). Are Emotions Natural Kinds? Perspectives on Psychological Science, 1(1), 28–58. https://doi.org/10.1111/j.1745-6916.2006.00003.x

Barrett, L. (2017). How emotions are made: The secret life of the brain. New York: Houghton Mifflin Harcourt.

Barrett, L., Mesquita, B., & Gendron, M. (2011). Context in Emotion Perception. Current Directions in Psychological Science, 20(5), 286–290. https://doi.org/10.1177/0963721411422522

Barrett, L., & Satpute, A. (2013). Large-scale brain networks in affective and social neuroscience: towards an integrative functional architecture of the brain. Current Opinion in Neurobiology, 23(3), 361–372. https://doi.org/10.1016/j.conb.2012.12.012

Barrett, L., & Simmons, W. (2015). Interoceptive predictions in the brain. Nature Reviews. Neuroscience, 16(7), 419–429. https://doi.org/10.1038/nrn3950

Baslet, G., Termini, L., & Herbener, E. (2009). Deficits in emotional awareness in schizophrenia and their relationship with other measures of functioning. The Journal of Nervous and Mental Disease, 197(9), 655.

Bastos, A., Usrey, W., Adams, R., Mangun, G., Fries, P., & Friston, K. (2012). Canonical microcircuits for predictive coding. Neuron, 76(4), 695–711. https://doi.org/10.1016/j.neuron.2012.10.038

Berthoz, S., Ouhayoun, B., & Parage, N. (2000). Etude preliminaire des niveaux de conscience emotionnelle chez des patients deprimes et des controles. (Preliminary study of the levels of emotional awareness in depressed patients and controls.). Annals of Medical Psychology (Paris), 158, 665–672.

Binder, J., Desai, R., Graves, W., & Conant, L. (2009). Where Is the Semantic System? A Critical Review and Meta-Analysis of 120 Functional Neuroimaging Studies. Cerebral Cortex, 19(12), 2767–2796. https://doi.org/10.1093/cercor/bhp055

Bréjard, V., Bonnet, A., & Pedinielli, J. (2012). The role of temperament and emotional awareness in risk taking in adolescents. L’Encéphale: Revue de Psychiatrie Clinique Biologique et Thérapeutique, 38(1), 1–9.

Brown, T. H., Zhao, Y., & Leung, V. (2010). Hebbian plasticity. In Encyclopedia of Neuroscience (pp. 1049–1056). https://doi.org/10.1016/B978-008045046-9.00796-8

Burger, A., Lumley, M., Carty, J., Latsch, D., Thakur, E., Hyde-Nolan, M., … Schubiner, H. (2016). The effects of a novel psychological attribution and emotional awareness and expression therapy for chronic musculoskeletal pain: A preliminary, uncontrolled trial. Journal of Psychosomatic Research, 81, 1–8. https://doi.org/10.1016/j.jpsychores.2015.12.003

Bydlowski, S., Corcos, M., Jeammet, P., Paterniti, S., Berthoz, S., Laurier, C., … Consoli, S. (2005). Emotion-processing deficits in eating disorders. International Journal of Eating Disorders, 37, 321–329.

Chancey, J., Adlaf, E., Sapp, M., Pugh, P., Wadiche, J., & Overstreet-Wadiche, L. (2013). GABA depolarization is required for experience-dependent synapse unsilencing in adult-born neurons. The Journal of Neuroscience : The Official Journal of the Society for Neuroscience, 33(15), 6614–6622. https://doi.org/10.1523/JNEUROSCI.0781-13.2013

Chaplin, T., Cole, P., & Zahn-Waxler, C. (2005). Parental Socialization of Emotion Expression: Gender Differences and Relations to Child Adjustment. Emotion, 5(1), 80–88. https://doi.org/10.1037/1528-3542.5.1.80

Ciarrochi, J., Caputi, P., & Mayer, J. (2003). The distinctiveness and utility of a measure of trait emotional awareness. Personality and Individual Differences, 34, 1477–1490.

Clark, J., Watson, S., & Friston, K. (2018). What is mood? A computational perspective. Psychological Medicine, 1–8. https://doi.org/10.1017/S0033291718000430

Colvert, E., Rutter, M., Kreppner, J., Beckett, C., Castle, J., Groothues, C., … Sonuga-Barke, E. (2008). Do Theory of Mind and Executive Function Deficits Underlie the Adverse Outcomes Associated with Profound Early Deprivation?: Findings from the English and Romanian Adoptees Study. Journal of Abnormal Child Psychology, 36(7), 1057–1068. https://doi.org/10.1007/s10802-008-9232-x

Conant, C., & Ashbey, W. (1970). Every good regulator of a system must be a model of that system. International Journal of Systems Science, 1(2), 89–97. https://doi.org/10.1080/00207727008920220

Consoli, S., Lemogne, C., Roch, B., Laurent, S., Plouin, P., & Lane, R. (2010). Differences in emotion processing in patients with essential and secondary hypertension. American Journal of Hypertension, 23, 515–521.

Cooper, A. A., Clifton, E. G., & Feeny, N. C. (2017, August 1). An empirical review of potential mediators and mechanisms of prolonged exposure therapy. Clinical Psychology Review, Vol. 56, pp. 106–121. https://doi.org/10.1016/j.cpr.2017.07.003

Darby, K., & Sloutsky, V. (2015). The cost of learning: interference effects in memory development. Journal of Experimental Psychology. General, 144(2), 410–431. https://doi.org/10.1037/xge0000051

de Berker, A., Rutledge, R., Mathys, C., Marshall, L., Cross, G., Dolan, R., & Bestmann, S. (2016). Computations of uncertainty mediate acute stress responses in humans. Nature Communications, 7, 10996. https://doi.org/10.1038/ncomms10996

Diener, M., & Lucas, R. (2004). Adults Desires for Childrens Emotions across 48 Countries: Associations with Individual and National Characteristics. Journal of Cross-Cultural Psychology, 35(5), 525–547. https://doi.org/10.1177/0022022104268387

Donges, U., Kersting, A., Dannlowski, U., Lalee-Mentzel, J., Arolt, V., & Suslow, T. (2005). Reduced awareness of others’ emotions in unipolar depressed patients. The Journal of Nervous and Mental Disease, 193(5), 331–337.

Erbas, Y., Ceulemans, E., Boonen, J., Noens, I., & Kuppens, P. (2013). Emotion differentiation in autism spectrum disorder. Research in Autism Spectrum Disorders, 7(10), 1221–1227. https://doi.org/10.1016/j.rasd.2013.07.007

Fivush, R., Brotman, M., Buckner, J., & Goodman, S. (2000). Gender Differences in Parent–Child Emotion Narratives. Sex Roles, 42(3), 233–253. https://doi.org/10.1023/A:1007091207068

Frewen, P., Lane, R., Neufeld, R., Densmore, M., Stevens, T., & Lanius, R. (2008). Neural correlates of levels of emotional awareness during trauma script-imagery in posttraumatic stress disorder. Psychosomatic Medicine, 70(1), 27– 31.

Friston, K., FitzGerald, T., Rigoli, F., Schwartenbeck, P., O Doherty, J., & Pezzulo, G. (2016). Active inference and learning. Neuroscience and Biobehavioral Reviews, 68, 862–879. https://doi.org/10.1016/j.neubiorev.2016.06.022

Friston, K., FitzGerald, T., Rigoli, F., Schwartenbeck, P., & Pezzulo, G. (2017). Active Inference: A Process Theory. Neural Computation, 29(1), 1–49. https://doi.org/10.1162/NECO_a_00912

Friston, K., Lin, M., Frith, C., Pezzulo, G., Hobson, J., & Ondobaka, S. (2017). Active Inference, Curiosity and Insight. Neural Computation, 29(10), 2633–2683. https://doi.org/10.1162/neco_a_00999

Friston, K., Parr, T., & de Vries, B. (2017). The graphical brain: Belief propagation and active inference. Network Neuroscience, 1(4), 381–414. https://doi.org/10.1162/NETN_a_00018

Friston, K., Rosch, R., Parr, T., Price, C., & Bowman, H. (2018). Deep temporal models and active inference. Neuroscience & Biobehavioral Reviews, 90, 486–501. https://doi.org/10.1016/J.NEUBIOREV.2018.04.004

Funahashi, R., Maruyama, T., Yoshimura, Y., & Komatsu, Y. (2013). Silent synapses persist into adulthood in layer 2/3 pyramidal neurons of visual cortex in dark-reared mice. Journal of Neurophysiology, 109(8), 2064–2076. https://doi.org/10.1152/jn.00912.2012

Gusnard, D., Akbudak, E., Shulman, G., & Raichle, M. (2001). Medial prefrontal cortex and self-referential mental activity: relation to a default mode of brain function. Proceedings of the National Academy of Sciences, 98(7), 4259–4264.

Haker, H., Schneebeli, M., & Stephan, K. (2016). Can Bayesian Theories of Autism Spectrum Disorder Help Improve Clinical Practice? Frontiers in Psychiatry, 7, 107. https://doi.org/10.3389/fpsyt.2016.00107

Harmon-Jones, E., Gable, P., & Peterson, C. (2010). The role of asymmetric frontal cortical activity in emotion-related phenomena: A review and update. Biological Psychology, 84(3), 451–462. https://doi.org/10.1016/J.BIOPSYCHO.2009.08.010

Hayes, S., & Smith, S. (2005). Get Out of Your Mind and Into Your Life: The New Acceptance and Commitment Therapy. Oakland, CA: New Harbinger Publications.

Hietanen, J., Glerean, E., Hari, R., & Nummenmaa, L. (2016). Bodily maps of emotions across child development. Developmental Science, 19(6), 1111–1118. https://doi.org/10.1111/desc.12389

Hobson, J., & Friston, K. (2012). Waking and dreaming consciousness: neurobiological and functional considerations. Progress in Neurobiology, 98(1), 82–98. https://doi.org/10.1016/j.pneurobio.2012.05.003

Hohwy, J. (2016). The Self-Evidencing Brain. Noûs, 50(2), 259–285. https://doi.org/10.1111/nous.12062

Joffily, M., & Coricelli, G. (2013). Emotional Valence and the Free-Energy Principle. PLoS Computational Biology, 9(6), e1003094. https://doi.org/10.1371/journal.pcbi.1003094

Kashdan, T., Barrett, L., & McKnight, P. (2015). Unpacking Emotion Differentiation: Transforming Unpleasant Experience by Perceiving Distinctions in Negativity. Current Directions in Psychological Science, 24(1), 10–16. https://doi.org/10.1177/0963721414550708

Kashdan, T., & Farmer, A. (2014). Differentiating emotions across contexts: comparing adults with and without social anxiety disorder using random, social interaction, and daily experience sampling. Emotion, 14(3), 629–638. https://doi.org/10.1037/a0035796

Kerchner, G., & Nicoll, R. (2008). Silent synapses and the emergence of a postsynaptic mechanism for LTP. Nature Reviews. Neuroscience, 9(11), 813– 825. https://doi.org/10.1038/nrn2501

Kleckner, I., Zhang, J., Touroutoglou, A., Chanes, L., Xia, C., Simmons, W., … Barrett, L. (2017). Evidence for a Large-Scale Brain System Supporting Allostasis and Interoception in Humans. Nature Human Behaviour, 1, 0069. https://doi.org/10.1038/s41562-017-0069

Lackner, J. (2005). Is IBS a problem of emotion dysregulation? Testing the levels of emotional awareness model. Presented at the Annual Meeting of the American Psychosomatic Society.

Lane, R., Anderson, F., & Smith, R. (2018). Biased Competition Favoring Physical Over Emotional Pain: A Possible Explanation for the Link Between Early Adversity and Chronic Pain. Psychosomatic Medicine, 80(9), 880–890. https://doi.org/10.1097/PSY.0000000000000640

Lane, R., Fink, G., Chua, P., & Dolan, R. (1997). Neural activation during selective attention to subjective emotional responses. Neuroreport, 8(18), 3969–3972.

Lane, R., Quinlan, D., Schwartz, G., Walker, P., & Zeitlin, S. (1990). The Levels of Emotional Awareness Scale: a cognitive-developmental measure of emotion. Journal of Personality Assessment, 55(1–2), 124–134. https://doi.org/10.1080/00223891.1990.9674052

Lane, R., & Schwartz, G. (1987). Levels of emotional awareness: a cognitive-developmental theory and its application to psychopathology. American Journal of Psychiatry, 144, 133–143.

Lane, R., Sechrest, L., Reidel, R., Weldon, V., Kaszniak, A., & Schwartz, G. (1996). Impaired verbal and nonverbal emotion recognition in alexithymia. Psychosomatic Medicine, 58(3), 203–210.

Lane, R., Sechrest, L., Riedel, R., Shapiro, D., & Kaszniak, A. (2000). Pervasive emotion recognition deficit common to alexithymia and the repressive coping style. Psychosomatic Medicine, 62, 492–501.

Lane, R., Weihs, K., Herring, A., Hishaw, A., & Smith, R. (2015). Affective agnosia: Expansion of the alexithymia construct and a new opportunity to integrate and extend Freud’s legacy. Neuroscience and Biobehavioral Reviews, 55, 594–611. https://doi.org/10.1016/j.neubiorev.2015.06.007

Lawson, R., Mathys, C., & Rees, G. (2017). Adults with autism overestimate the volatility of the sensory environment. Nature Neuroscience, 20(9), 1293–1299. https://doi.org/10.1038/nn.4615

Lawson, R., Rees, G., & Friston, K. (2014). An aberrant precision account of autism. Frontiers in Human Neuroscience, 8, 302. https://doi.org/10.3389/fnhum.2014.00302

Levine, D., Marziali, E., & Hood, J. (1997). Emotion processing in borderline personality disorders. Journal of Nervous and Mental Disease, 185, 240–246.

Licata, M., Kristen, S., & Sodian, B. (2016). Mother-Child Interaction as a Cradle of Theory of Mind: The Role of Maternal Emotional Availability. Social Development, 25(1), 139–156. https://doi.org/10.1111/sode.12131

Limanowski, J., & Friston, K. (2018). “Seeing the Dark”: Grounding Phenomenal Transparency and Opacity in Precision Estimation for Active Inference. Frontiers in Psychology, 9, 643. https://doi.org/10.3389/fpsyg.2018.00643

Lindquist, K., & Barrett, L. (2008). Constructing emotion: the experience of fear as a conceptual act. Psychological Science, 19(9), 898–903. https://doi.org/10.1111/j.1467-9280.2008.02174.x

Lumley, M., Schubiner, H., Lockhart, N., Kidwell, K., Harte, S., Clauw, D., & Williams, D. (2017). Emotional awareness and expression therapy, cognitive behavioral therapy, and education for fibromyalgia: a cluster-randomized controlled trial. Pain, 158(12), 2354–2363. https://doi.org/10.1097/j.pain.0000000000001036

Martínez, M., Villar, M. E., Ballarini, F., & Viola, H. (2014). Retroactive interference of object-in-context long-term memory: Role of dorsal hippocampus and medial prefrontal cortex. Hippocampus, 24(12), 1482–1492. https://doi.org/10.1002/hipo.22328

McKay, R., & Dennett, D. (2009). The evolution of misbelief. Behavioral and Brain Sciences, 32(06), 493. https://doi.org/10.1017/S0140525X09990975

McRae, K., Reiman, E., Fort, C., Chen, K., & Lane, R. (2008). Association between trait emotional awareness and dorsal anterior cingulate activity during emotion is arousal-dependent. NeuroImage, 41(2), 648–655. https://doi.org/10.1016/j.neuroimage.2008.02.030

Metzinger, T. (2017). The Problem of Mental Action. In T. Metzinger & W. Wiese. (Eds.), Philosophy and Predicitive Processing.

Mirza, M., Adams, R., Mathys, C., & Friston, K. (2016). Scene Construction, Visual Foraging, and Active Inference. Frontiers in Computational Neuroscience, 10, 56. https://doi.org/10.3389/fncom.2016.00056

Moeller, S., Konova, A., Parvaz, M., Tomasi, D., Lane, R., Fort, C., … RZ, G. (2014). Functional, Structural, and Emotional Correlates of Impaired Insight in Cocaine Addiction. JAMA Psychiatry, 71(1), 61. https://doi.org/10.1001/jamapsychiatry.2013.2833

Mogg, K., & Bradley, B. (2005). Attentional Bias in Generalized Anxiety Disorder Versus Depressive Disorder. Cognitive Therapy and Research, 29(1), 29–45. https://doi.org/10.1007/s10608-005-1646-y

Mumme, D., Fernald, A., & Herrera, C. (1996). Infants’ Responses to Facial and Vocal Emotional Signals in a Social Referencing Paradigm. Child Development, 67(6), 3219–3237. https://doi.org/10.1111/j.1467-8624.1996.tb01910.x

Panksepp, J., & Biven, L. (2012). The Archaeology of Mind: Neuroevolutionary Origins of Human Emotions. New York: W.W. Norton & Company.

Panksepp, J., Lane, R., Solms, M., & Smith, R. (2017). Reconciling cognitive and affective neuroscience perspectives on the brain basis of emotional experience. Neuroscience and Biobehavioral Reviews, 76, *part B*, 187–215. https://doi.org/10.1016/j.neubiorev.2016.09.010

Parker, J., Taylor, G., & Bagby, R. (2003). The 20-Item Toronto Alexithymia Scale: III. Reliability and factorial validity in a community population. Journal of Psychosomatic Research, 55(3), 269–275. https://doi.org/10.1016/S0022-3999(02)00578-0

Parr, T., & Friston, K. (2017a). Uncertainty, epistemics and active inference. Journal of the Royal Society, Interface, 14(136). https://doi.org/10.1098/rsif.2017.0376

Parr, T., & Friston, K. (2017b). Working memory, attention, and salience in active inference. Scientific Reports, 7(1), 14678. https://doi.org/10.1038/s41598-017-15249-0

Parr, T., & Friston, K. (2018). The Anatomy of Inference: Generative Models and Brain Structure. Frontiers in Computational Neuroscience, 12, 90. https://doi.org/10.3389/fncom.2018.00090

Pears, K., & Fisher, P. (2005). Emotion understanding and theory of mind among maltreated children in foster care: Evidence of deficits. Development and Psychopathology, 17(01), 47–65. https://doi.org/10.1017/S0954579405050030

Peters, A., McEwen, B. S., & Friston, K. (2017). Uncertainty and stress: Why it causes diseases and how it is mastered by the brain. Progress in Neurobiology, 156, 164–188. https://doi.org/10.1016/J.PNEUROBIO.2017.05.004

Posner, M. (2016). Orienting of attention: Then and now. Quarterly Journal of Experimental Psychology (2006), 69(10), 1864–1875. https://doi.org/10.1080/17470218.2014.937446

Rizzolatti, G., Riggio, L., Dascola, I., & Umiltá, C. (1987). Reorienting attention across the horizontal and vertical meridians: Evidence in favor of a premotor theory of attention. Neuropsychologia, 25(1), 31–40. https://doi.org/10.1016/0028-3932(87)90041-8

Russell, J. (1991). Culture and the categorization of emotions. Psychological Bulletin, 110(3), 426–450. https://doi.org/10.1037/0033-2909.110.3.426

Russell, J. (2003). Core affect and the psychological construction of emotion. Psychological Review, 110(1), 145–172.

Scherer, K. (2009). The dynamic architecture of emotion: Evidence for the component process model. Cognition & Emotion, 23(7), 1307–1351. https://doi.org/10.1080/02699930902928969

Schwartenbeck, P., FitzGerald, T., Mathys, C., Dolan, R., & Friston, K. (2015). The Dopaminergic Midbrain Encodes the Expected Certainty about Desired Outcomes. Cerebral Cortex, 25(10), 3434–3445. https://doi.org/10.1093/cercor/bhu159

Schwartenbeck, P., & Friston, K. (2016). Computational Phenotyping in Psychiatry: A Worked Example. ENeuro, 3(4), ENEURO.0049-0016.2016. https://doi.org/10.1523/ENEURO.0049-16.2016

Seth, A. (2013). Interoceptive inference, emotion, and the embodied self. Trends in Cognitive Sciences, 17(11), 565–573. https://doi.org/10.1016/j.tics.2013.09.007

Seth, A., & Friston, K. (2016). Active interoceptive inference and the emotional brain. Philosophical Transactions of the Royal Society of London B: Biological Sciences, 371(1708).

Sharot, T. (2011). The optimism bias. Current Biology, 21(23), R941–R945. https://doi.org/10.1016/J.CUB.2011.10.030

Siemer, M., Mauss, I., & Gross, J. (2007). Same situation--Different emotions: How appraisals shape our emotions. Emotion, 7(3), 592–600. https://doi.org/10.1037/1528-3542.7.3.592

Silani, G., Bird, G., Brindley, R., Singer, T., Frith, C., & Frith, U. (2008). Levels of emotional awareness and autism : An fMRI study Levels of emotional awareness and autism : An fMRI study. Psychology, 3(2), 97–112. https://doi.org/10.1080/17470910701577020

Smith, D., & Schenk, T. (2012). The Premotor theory of attention: Time to move on? Neuropsychologia, 50(6), 1104–1114. https://doi.org/10.1016/J.NEUROPSYCHOLOGIA.2012.01.025

Smith, R. (2019). The three-process model of implicit and explicit emotion. In R. Lane & L. Nadel (Eds.), Neuroscience of Enduring Change: Implications for Psychotherapy (p. (in press)). Oxford University Press.

Smith, R., Akozei, A., Killgore, W., & Lane, R. (2017). Nested Positive Feedback Loops in the Maintenance of Major Depression: An integration and extension of previous models. Brain, Behavior, and Immunity, (In Press). https://doi.org/10.1016/j.bbi.2017.09.011

Smith, R., Alkozei, A., Bao, J., Smith, C., Lane, R., & Killgore, W. (2017). Resting state functional connectivity correlates of emotional awareness. NeuroImage, 159, 99–106. https://doi.org/10.1016/j.neuroimage.2017.07.044

Smith, R., Bajaj, S., Dailey, N., Alkozei, A., Smith, C., Sanova, A., … Killgore, W. (2018). Greater cortical thickness within the limbic visceromotor network predicts higher levels of trait emotional awareness. Consciousness and Cognition, 57, 54– 61.

Smith, R., Braden, B., Chen, K., Ponce, F., Lane, R., & Baxter, L. (2015). The neural basis of attaining conscious awareness of sad mood. Brain Imaging and Behavior, 9(3), 574–587. https://doi.org/10.1007/s11682-014-9318-8

Smith, R., Fass, H., & Lane, R. (2014). Role of medial prefrontal cortex in representing one’s own subjective emotional responses: A preliminary study. Consciousness and Cognition, 29, 117–130. https://doi.org/10.1016/j.concog.2014.08.002

Smith, R., Kaszniak, A., Katsanis, J., Lane, R., & Nielsen, L. (2019). The importance of identifying underlying process abnormalities in alexithymia: Implications of the three-process model and a single case study illustration. Consciousness and Cognition, 68, 33–46. https://doi.org/10.1016/J.CONCOG.2018.12.004

Smith, R., Killgore, W., & Lane, R. (2018). The structure of emotional experience and its relation to trait emotional awareness: A theoretical review. Emotion, 18(5), 670–692. https://doi.org/10.1037/emo0000376

Smith, R., & Lane, R. (2015). The neural basis of one’s own conscious and unconscious emotional states. Neuroscience and Biobehavioral Reviews, 57, 1– 29. https://doi.org/10.1016/j.neubiorev.2015.08.003

Smith, R., & Lane, R. (2016). Unconscious emotion: A cognitive neuroscientific perspective. Neuroscience and Biobehavioral Reviews, 69, 216–238. https://doi.org/10.1016/j.neubiorev.2016.08.013

Smith, R., Lane, R., Alkozei, A., Bao, J., Smith, C., Sanova, A., … Killgore, W. (2017). Maintaining the feelings of others in working memory is associated with activation of the left anterior insula and left frontal-parietal control network. Social Cognitive and Affective Neuroscience, 12(5), 848–860. https://doi.org/10.1093/scan/nsx011

Smith, R., Lane, R., Alkozei, A., Bao, J., Smith, C., Sanova, A., … Killgore, W. (2018). The role of medial prefrontal cortex in the working memory maintenance of one’s own emotional responses. Scientific Reports, 8. https://doi.org/10.1038/s41598-018-21896-8

Smith, R., Lane, R., Parr, T., & Friston, K. (2019). Neurocomputational mechanisms underlying emotional awareness: insights afforded by deep active inference and their potential clinical relevance. Neuroscience and Biobehavioral Reviews. https://doi.org/10.1101/681288

Smith, R., Lane, R., Sanova, A., Alkozei, A., Smith, C., & Killgore, W. W. D. (2018). Common and Unique Neural Systems Underlying the Working Memory Maintenance of Emotional vs. Bodily Reactions to Affective Stimuli: The Moderating Role of Trait Emotional Awareness. Frontiers in Human Neuroscience, 12, 370. https://doi.org/10.3389/fnhum.2018.00370

Smith, R., Quinlan, D., Schwartz, G. E., Sanova, A., Alkozei, A., & Lane, R. D. (2019). Developmental Contributions to Emotional Awareness. Journal of Personality Assessment, 101(2), 150–158. https://doi.org/10.1080/00223891.2017.1411917

Smith, R., Sanova, A., Alkozei, A., Lane, R., & Killgore, W. (2018). Higher levels of trait emotional awareness are associated with more efficient global information integration throughout the brain: a graph-theoretic analysis of resting state functional connectivity. Social Cognitive and Affective Neuroscience, 13(7), 665– 675. https://doi.org/10.1093/scan/nsy047

Smith, R., Schwartenbeck, P., Parr, T., & Friston, K. (2019). An active inference approach to modeling structure learning: concept learning as an example case. BioRxiv, 633677. https://doi.org/10.1101/633677

Smith, R., Thayer, J., Khalsa, S., & Lane, R. (2017). The hierarchical basis of neurovisceral integration. Neuroscience and Biobehavioral Reviews, 75, 274– 296. https://doi.org/10.1016/j.neubiorev.2017.02.003

Smith, R., Weihs, K., Alkozei, A., Killgore, W., & Lane, R. (2019). An embodied neurocomputational framework for organically integrating biopsychosocial processes: An application to the role of social support in health and disease. Psychosomatic Medicine. https://doi.org/10.1097/PSY.0000000000000661

Subic-Wrana, A., Beetz, M., Paulussen, J., Wiltnik, J., & Beutel, M. (2007). Relations between attachment, childhood trauma, and emotional awareness in psychosomatic inpatients. Budapest, Hungary.

Subic-Wrana, C., Bruder, S., Thomas, W., Lane, R., & Köhle, K. (2005). Emotional awareness deficits in inpatients of a psychosomatic ward: a comparison of two different measures of alexithymia. Psychosomatic Medicine, 67(3), 483–489.

Suvak, M., Litz, B., Sloan, D., Zanarini, M., Barrett, L., & Hofmann, S. (2011). Emotional granularity and borderline personality disorder. Journal of Abnormal Psychology, 120(2), 414–426. https://doi.org/10.1037/a0021808

Tononi, G., & Cirelli, C. (2014). Sleep and the Price of Plasticity: From Synaptic and Cellular Homeostasis to Memory Consolidation and Integration. Neuron, 81(1), 12–34. https://doi.org/10.1016/J.NEURON.2013.12.025

Widen, S., & Russell, J. (2008). Children acquire emotion categories gradually. Cognitive Development, 23(2), 291–312. https://doi.org/10.1016/j.cogdev.2008.01.002

Wilson-Mendenhall, C., Barrett, L., Simmons, W., & Barsalou, L. (2011). Grounding emotion in situated conceptualization. Neuropsychologia, 49(5), 1105–1127. https://doi.org/10.1016/j.neuropsychologia.2010.12.032

Wright, R., Riedel, R., Sechrest, L., Lane, R., & Smith, R. (2017). Sex differences in emotion recognition ability: The mediating role of trait emotional awareness. Motivation and Emotion. https://doi.org/10.1007/s11031-017-9648-0

Yeo, B., Krienen, F., Sepulcre, J., Sabuncu, M., Lashkari, D., Hollinshead, M., … Buckner, R. (2011). The organization of the human cerebral cortex estimated by intrinsic functional connectivity. Journal of Neurophysiology, 106(3), 1125– 1165. https://doi.org/10.1152/jn.00338.2011

