## Supplementary material for "Simulating emotions: An active inference model of emotional state inference and emotion concept learning": Emotion_learning_model.m

```
%% Set up emotion inference model%__________________________________________________________________________
clear
close all
rng('shuffle')

% Simulation options
%--------------------------------------------------------------------------
trial_n = 1; % number of trials: 1 = single trial, 2 = 200 trials

learn_a = 0; % set to 1 to allow emotion concept learning (A-matrix)
learn_b = 0; % set to 1 to allow transition learning (B-matrix)
learn_d = 0; % set to 1 to allow learning of priors over emotional states (D-matrix)

attention_bias = 0; % 1 = internal bias, 2 = external bias, 3 = somatic bias

emotion = 1; % Choose initial emotional state: 1 = sad, 2 = afraid, 3 = angry, 4 = happy

% Precision (inverse temperature) parameters
%--------------------------------------------------------------------------

p_A_L1 = 2; % Attenuate A-matrix precision (Generative process)
p_A1_L1 = 2;% Attenuate the specific mapping from emotion concepts to observations (Generative process)
p_a_L1 = 0; % Attenuate the specific mapping from emotion concepts to observations (Generative model; if learning)
p_B_L1 = 2; % Attenuate B-matrix (transition) precision (generative process)
p_b_L1 = 0; % Attenuate B-matrix (transition) precision (generative model; if learning)

% prior beliefs about initial states (in terms of counts_: D and d
%--------------------------------------------------------------------------

% Generative process
D{1} = [1 1 1 1]';           % Emotion:     {'Sad','Afraid','Angry', 'Happy'}
D{2} = [1 0 0 0 0 0 0 0 0 0]'; % Attention Allocation:    {'start','Valence','Arousal','Action Tendency','Beliefs','Context',
                                                          %'Choose Sad','Choose Afreaid','Choose Angry','Choose Happy'}
% Generative model (if learning)
d{1} = [1 1 1 1]';
d{2} = D{2};

% probabilistic mapping from hidden states to outcomes: A
%--------------------------------------------------------------------------
Nf    = numel(D);
for f = 1:Nf
    Ns(f) = numel(D{f});
end
No    = [13 10]; % Lower-level representations, Attended Modality and feedback
Ng    = numel(No);
for g = 1:Ng
    A{g} = zeros([No(g),Ns]);
end


%A-Matrix: Mapping States to Observations
%--------------------------------------------------------------------------

% Mapping to lower-level representations
A{1}(1,:,1) = [1 1 1 1]; %('Attention to Start')

A{1}(2:3,:,2) = [0 0 0 1;  % Pleasant
                 1 1 1 0]; % Unpleasant
A{1}(4:5,:,3) = [1/2 0 0 1/2;  % Low arousal
                 1/2 1 1 1/2]; % High arousal
A{1}(6:7,:,4) = [0 0 1 1/2;  % Approach
                 1 1 0 1/2]; % Avoid
A{1}(8:9,:,5) = [1 0 0 1/2;  % Self-blame
                 0 1 1 1/2]; % Other-blame
A{1}(10:11,:,6) = [1 1/2 1 0;  % Social rejection
                   0 1/2 0 1]; % Crowded event

% Feedback: correct/incorrect
A{1}(12:13,:,7) = [1 0 0 0;0 1 1 1]; %('Choose Sad')
A{1}(12:13,:,8) = [0 1 0 0;1 0 1 1]; %('Choose Afriad')
A{1}(12:13,:,9) = [0 0 1 0;1 1 0 1]; %('Choose Angry')
A{1}(12:13,:,10)= [0 0 0 1;1 1 1 0]; %('Choose Happy')

% Mapping to attended modality
for i = 1:10
    A{2}(i,:,i) = [1 1 1 1];
end

for g = 1:Ng
    A{g} = double(A{g});
end

a = A; % Initially match generative model to generative process (A-matrix)

for i=1:length(A)
    A{i}=spm_softmax(p_A_L1*log(A{i}+exp(-4))); % Attenuate A-matrix (Generative process)
end

A{1}(:,:,2:6)=spm_softmax(p_A1_L1*log(A{1}(:,:,2:6)+exp(-4))); % Attenuate emotion concept knowledge (Generative process)

a{1}(:,:,2:6)=spm_softmax(p_a_L1*log(a{1}(:,:,2:6)+exp(-4))); % Attenuate emotion concept knowledge (Generative Model)


% B-matrix: State transition probabilities
%--------------------------------------------------------------------------
for f = 1:Nf
    B{f} = eye(Ns(f)); % State transitions between emotions are not controllable
end

for k = 1:Ns(2)
    B{2}(:,:,k) = 0;
    B{2}(k,:,k) = 1;
end

B{2}(:,7:10,:) = 0;

for i = 7:10
    B{2}(i,i,:) = 1; % State transitions between attentional states are controllable
end

b = B; % Initially match generative model to generative process (B-matrix)

B{1}=spm_softmax(p_B_L1*log(B{1}+exp(-4))); % Attenuate the stability of emotional states (Generative process)

b{1}=spm_softmax(p_b_L1*log(b{1}+exp(-4))); % Attenuate the stability of emotional states (Generative model)

% allowable policies (here, specified as the next action) U
%--------------------------------------------------------------------------
T = 6;
Np        = size(B{2},3);
U         = ones(1,Np,Nf);
U(:,:,2)  = 1:Np;

% C-Matrix: prior preferences
%--------------------------------------------------------------------------
C{1}      = zeros(No(1),T);
C{2}      = zeros(No(2),T);
C{1}(12,:) =  2;                 % the agent prefers to be right
C{1}(13,:) = -4;                 % and not wrong
C{2}(1:6,T:end) = -4;            % make Delayed sampling costly

% E-Matrix: attention biases
%--------------------------------------------------------------------------

E_INT = [1 1 1 1 1/50 1/50 1 1 1 1]';    % only internal focus
E_EXT = [1 1/50 1/50 1/50 1 1 1 1 1 1]'; % only external focus
E_SOM = [1 1/50 1 1 1/50 1/50 1 1 1 1]'; % somatic

% MDP Structure - this will be used to generate arrays for multiple trials
%==========================================================================
mdp.T = T;                      % number of moves
mdp.U = U;                     % allowable shallow policies
mdp.A = A;                      % observation model
mdp.B = B;                      % transition probabilities
mdp.C = C;                      % preferred outcomes
mdp.D = D;                      % prior over initial states

if attention_bias == 1
    mdp.E = E_INT;                      %habits
elseif attention_bias == 2
    mdp.E = E_EXT;
elseif attention_bias == 3
    mdp.E = E_SOM;
end

if learn_a == 1
    mdp.a = a;                      % observation beliefs
end

if learn_b == 1
    mdp.b = b;                      % transition beliefs
end

if learn_d == 1
    mdp.d = d;                      % prior beliefs over initial states
end

mdp.s = [emotion 1]';                          % initial state


label.factor{1}   = 'Emotion concepts';   label.name{1}    = {'Sad','Afraid','Angry','Happy'};
label.factor{2}   = 'Attentional focus';     label.name{2}    = {'Start','Valence','Arousal','Action tendencies','Beliefs','Context','Report sad','Report afraid','Report angry','Report Happy'};
label.modality{1} = 'Lower-level representations';    label.outcome{1} = {'Start','Pleasant','Unpleasant','Low arousal','High arousal','Approach','Avoid','Self-blame','Other-blame','Social rejection','Crowded event','Correct','Incorrect'};
label.modality{2} = 'Attended information';  label.outcome{2} = {'Start','Valence','Arousal','Action tendencies','Beliefs','Context','Report sad','Report afraid','Report angry','Report Happy'};
label.action{2} = 'Attended information';  label.action{2} = {'Start','Valence','Arousal','Action tendencies','Beliefs','Context','Report sad','Report afraid','Report angry','Report Happy'};


mdp.label = label;

mdp.Aname = {'Lower-level representations','Attended Information'};
mdp.Bname = {'Emotion Concepts','Attention Focus'};
mdp.alpha = 128;
mdp.beta = 1;

mdp         = spm_MDP_check(mdp);
%% illustrate a single trial
%==========================================================================
MDP   = spm_MDP_VB_X(mdp);

% show belief updates (and behaviour)
%--------------------------------------------------------------------------
spm_figure('GetWin','initial trial: behavior'); clf
spm_MDP_VB_trial(MDP);
subplot(3,2,3)

% illustrate phase-precession and responses
%--------------------------------------------------------------------------
spm_figure('GetWin','initial trial: physiology'); clf
spm_MDP_VB_LFP(MDP,[],1);


if trial_n == 2
%% illustrate a sequence of trials
%==========================================================================

% true initial states – with context change at trial 12
%--------------------------------------------------------------------------
clear MDP

order = [1 2 3 4 1 2 3 4 1 2 3 4 1 2 3 4 1 2 3 4 1 2 3 4 1 2 3 4 1 2 3 4 1 2 3 4 1 2 3 4];

s(1,:) = [order order order order order]; % 200 trials (50 per emotion, interleaved)

N      = length(s(1,:));

s(2,:) = ceil(rand(1,N)*1);

for i = 1:N
   MDP(i)   = mdp;      % create structure array
   MDP(i).s = s(:,i);   % context
end

% Solve - an example sequence
%==========================================================================
MDP  = spm_MDP_VB_X(MDP);

% illustrate behavioural responses and neuronal correlates
%--------------------------------------------------------------------------
spm_figure('GetWin','trial sequence: behavior'); clf
spm_MDP_VB_game(MDP);

% illustrate phase-amplitude (theta-gamma) coupling
%--------------------------------------------------------------------------
spm_figure('GetWin','trial sequence: physiology'); clf
spm_MDP_VB_LFP(MDP(1:1:N));


% illustrate example trial after learning
%==========================================================================

% show belief updates (and behaviour)
%--------------------------------------------------------------------------
spm_figure('GetWin','example trial after learning: behavior'); clf
spm_MDP_VB_trial(MDP(N-3));
subplot(3,2,3)

% illustrate phase-precession and responses
%--------------------------------------------------------------------------
spm_figure('GetWin','example trial after learning: physiology'); clf
spm_MDP_VB_LFP(MDP(N-3),[],1);

% performance improvement over time during 200 learning trials (in bins of
% 10 or 5 for all emotions or each emotion, respectively
%--------------------------------------------------------------------------

for i=1:N
Out(i) = MDP(i).o(1,T);
State(i) = MDP(i).s(1,T);

    if MDP(i).o(1,T) == 12
        Acc(i)=1;
    else
        Acc(i)=0;
    end
end

percent_accuracy_total = ((sum(Acc))/N)*100

bin = 10;

for i = 1:N/(bin)
percent_accuracy_over_time(i) = sum(Acc((i*bin)-(bin-1):bin*i))/(bin)*100;
end

percent_accuracy_over_time

bin_emo = 5;

Sad_Acc=Acc(State==1);


for i = 1:length(Sad_Acc)/(bin_emo)
percent_accuracy_sad_over_time(i) = sum(Sad_Acc((i*bin_emo)-(bin_emo-1):bin_emo*i))/(bin_emo)*100;
end

percent_accuracy_sad_over_time

Fear_Acc=Acc(State==2);

for i = 1:length(Fear_Acc)/(bin_emo)
percent_accuracy_fear_over_time(i) = sum(Fear_Acc((i*bin_emo)-(bin_emo-1):bin_emo*i))/(bin_emo)*100;
end

percent_accuracy_fear_over_time

Anger_Acc=Acc(State==3);

for i = 1:length(Anger_Acc)/(bin_emo)
percent_accuracy_angry_over_time(i) = sum(Anger_Acc((i*bin_emo)-(bin_emo-1):bin_emo*i))/(bin_emo)*100;
end

percent_accuracy_angry_over_time

Happy_Acc=Acc(State==4);

for i = 1:length(Happy_Acc)/(bin_emo)
percent_accuracy_happy_over_time(i) = sum(Happy_Acc((i*bin_emo)-(bin_emo-1):bin_emo*i))/(bin_emo)*100;
end

percent_accuracy_happy_over_time

end
```
